# Induction of cellular totipotency by NACC1-driven feed-forward regulation

**DOI:** 10.1101/2025.10.14.682353

**Authors:** Thulaj Meharwade, Gilberto Duran-Bishop, Loïck Joumier, Sabin Dhakal, Mohammed Usama, Sanidhya Jagdish, Élie Lambert, Yacine Kherdjemil, François Robert, Mohan Malleshaiah

## Abstract

Mammalian embryo development begins with totipotent cells. Despite its significance, gene regulatory mechanisms controlling the totipotent cell stage are poorly understood. Based on single-cell proteomics and transcriptomics, we identified nucleus accumbens-associated 1 (NACC1) as a critical regulator of totipotent-like cell state in mouse embryonic stem cells. Using a combination of genomics approaches, we found that NACC1 directly binds to gene-regulatory regions and favours chromatin accessibility to induce the expression of totipotency and zygotic genome activation genes, as well as retrotransposons associated with totipotency. In parallel, NACC1-regulated retrotransposons further modulate the expression of proximal totipotency genes, forming a coherent feed-forward mechanism that regulates totipotent-like cells. Furthermore, NACC1 is crucial for the progression of embryogenesis beyond the totipotency stage. Thus, we uncover a genome-level NACC1-driven feed-forward gene regulatory mechanism that governs totipotent-like cells and has a crucial role during embryonic development.

## INTRODUCTION

Totipotency is observed at the initial stages of embryonic development and is conserved across mammalian preimplantation development (*1, 2*). Totipotent cells serve as the foundational cells for the entirety of developing embryos, through their ability to produce both embryonic and extraembryonic tissues (*3–5*). Consequently, normal establishment of totipotency is indispensable for the successful progression of embryos to both pre- and post-implantation stages. Despite its evolutionary conservation and critical significance in the initiation of mammalian life, the gene regulatory mechanisms that underpin totipotency remain elusive. A comprehensive understanding of these mechanisms is crucial for our understanding of developmental and stem cell biology, as well as for disease modelling.

In contrast to totipotency, pluripotency has been extensively studied through the utilization of embryonic stem cells (ESCs) and induced pluripotent stem cells (iPSCs), leading to the characterization of key gene regulatory factors and mechanisms governing it (*6–10*). In the absence of appropriate totipotent cell culturing systems, the plasticity of stem cells has been harnessed to reprogram ESCs into a totipotent-like cell (TLC) state (*11–20*). The TLCs embody mouse 2-Cell (2C) or human 8-Cell blastomeres by expressing signature genes of totipotency, such as *Dux*, *Zscan4*, and *Zfp352,* as well as zygotic genome activation (ZGA) genes (*11, 21–27*). The modelling of totipotency through ESCs has enabled the identification of several negative regulators of the mouse ESC-to-TLC transition, including LSD1/KDM1A, CAF-1, MYC, DNMT1, LIN28, ZMYM2, and miR-34a (*27–32*). However, positive regulators of totipotency remain scarce, with DUX, DPPA2/4, and NELFA being notable examples (*33–36*).

Specific gene expression programs distinguish the mouse TLCs and ESCs, characterized by the expression of totipotency genes (e.g., *Duxf3*, *Zscan4,* and *Zfp352*) and pluripotency genes (e.g., *Oct4* (*Pou5f1*), *Nanog*, *Klf4,* and *Sox2*), respectively (*11, 21, 22*). Additionally, a key characteristic of TLCs is the expression of endogenous retroviruses or retrotransposons (*11, 25*). Mouse TLCs express long terminal repeat (LTR) retrotransposons, such as murine endogenous retrovirus (MERVL) and MT2_Mm, which represent the two highly expressed retrotransposon families in the 2C blastomeres (*11, 37*). Retrotransposons occupy more than one-third of the mammalian genome, and their expression regulates preimplantation embryo development (*11, 37–39*). Despite their importance, only a few transcription factors (TF), including DUX and ZSCAN4c, have been demonstrated to directly bind and transcriptionally activate retrotransposons in mammalian TLCs (*32, 40–44*).

A coordinated function of developmental factors, in the form of regulatory networks, controls cell fate choice (*45–47*). Often, the evolutionarily conserved network motifs or circuits, such as feed-forward motifs, play a crucial role in regulating cell fate choices across species (*47–52*). In ESCs, pluripotency factors are observed to form extensive regulatory networks consisting of feed-forward, positive and negative feedback, and mutual inhibitory circuits to either maintain a stem cell state or induce differentiation (*53–57*). The structure of the networks that regulate TLCs remains largely unknown (Fig. 1A). Moreover, the involvement of retrotransposons in these networks, and whether such networks operate at the genome level, are poorly understood.

**Figure 1:**
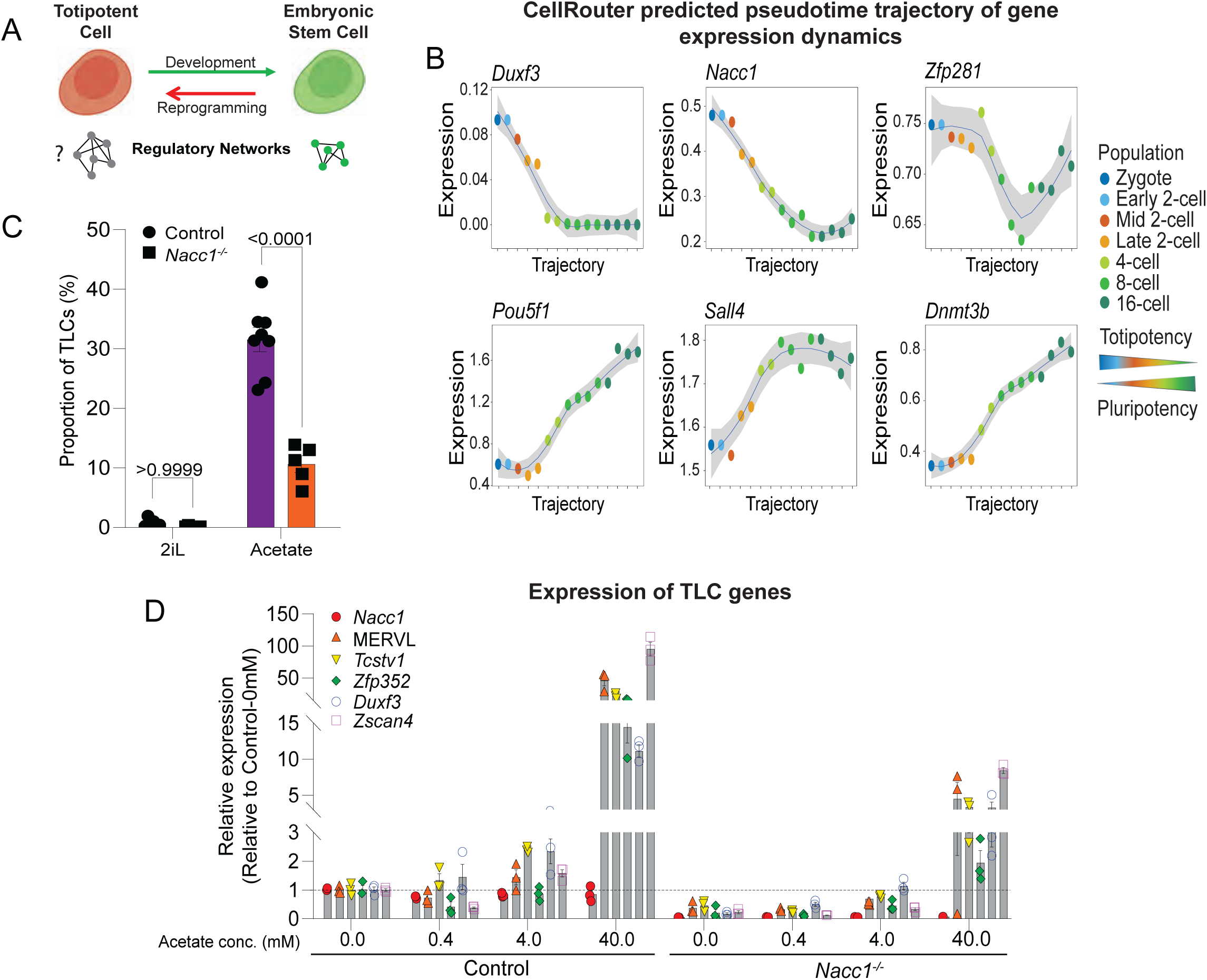
NACC1 is required for the TLC state in mouse ESCs. **(A)** Schematic showing the transition of totipotent cells to ESCs during development and culture-based reprogramming of ESCs to TLCs. Network schematics represent the well-characterized regulatory networks of ESCs and their poor understanding in totipotent cells. **(B)** Expression pattern of candidate transcription factors (*Duxf3*, *Nacc1*, *Zfp281*, *Pou5f1*, *Sall4*, *Dnmt3b*) along the CellRouter predicted pseudotime trajectory from zygote to 16-Cell stage of mouse embryogenesis. Blue line represents average expression while gray line represents standard deviation. **(C)** Proportion of TLCs (MERVL-gag positive cells) in *Nacc1*^-/-^ (*Nacc1*-KO gRNA2-Clone2) ESCs cultured in either 2iL or acetate conditions. **(D)** RT-qPCR analysis for expression of TLC genes in control and *Nacc1^-/-^* ESCs cultured with increasing concentration of acetate. All samples were normalized to control (SL + acetate (0 mM)). Measurements in C and D were done at 72h cultures. Graphs represent mean ± SEM in C and D. n=5-10 (in C) and n=3 (in D). *P* values were determined by Student’s t-test in (C).

In this study, we uncover a genome-level feed-forward gene regulatory mechanism that governs the TLCs and demonstrate its critical role in preimplantation embryo development. Central to this mechanism, we identify Nucleus Accumbens Associated 1 (NACC1) (*55, 58, 59*) as a crucial factor required for the transition of ESCs to the TLC state and for the progression of embryos beyond the totipotency stage. At the molecular level, NACC1 induces the expression of totipotency genes both directly and through the activation of retrotransposon expression. These findings reveal a potentially conserved feed-forward gene regulatory mechanism underlying the totipotent state and the initiation of mammalian embryo development.

## RESULTS

### NACC1 is crucial for ESC transition to the TLC state

Pluripotency and differentiation of mouse ESCs are regulated by multiple developmental TFs (*6, 54, 55, 60, 61*). Given the pivotal role of TFs in cell fate specification, we hypothesized that at least a subset of TFs regulating ESCs might also contribute to the establishment of TLCs (Fig. 1A). To identify such TLC TFs, we compared profiles of multiple TFs previously quantified by single-cell mass cytometry across ESC cell states (*62*). TLC and pluripotent cells were identified using ZSCAN4 and POU5F1 (OCT4) protein level, respectively (*62*). While the median protein levels for most TFs were high in pluripotent cells, a subset - including SALL4, NACC1, and ZFP281-also showed high expression levels in TLCs (fig. S1A-B). A correlation network analysis suggested potential links between ZSCAN4 (a TLC marker), SALL4, NACC1, and other pluripotency TFs such as POU5F1 and SOX2 (fig. S1C). Given that the core pluripotency TFs, POU5F1, SOX2, NANOG and KLF4 protein levels were relatively low and shown to be suppressed in TLCs, we focused on the other pluripotency-associated proteins, namely SALL4, NACC1, ZFP281 and DNMT3B (fig. S1A) (*12, 28*).

To further assess the expression dynamics of *Sall4*, *Nacc1*, *Zfp281* and *Dnmt3b* genes during preimplantation, we performed CellRouter trajectory analysis of available single-cell mRNA sequencing (scRNA-seq) data (*63–65*). The pseudotime trajectory, spanning from the Zygote to the 16-cell stage, revealed a contrasting expression pattern between TLC and pluripotency genes (Fig. 1B and S1D). While the expression of TLC genes, such as *Duxf3* and *Zscan4a-f,* was high at the zygote and 2-Cell stages and declined sharply by the 16-Cell stage, an inverse pattern is observed for pluripotency genes, including *Pou5f1*, *Nanog*, *Tcfcp2l1* and *Esrrb* (Fig. 1B and S1D). Among the potential new TLC genes, only *Nacc1* mirrored the *Duxf3* expression pattern, *Zfp281* transiently decreased at 4 and 8-cell stages, and *Sall4* and *Dnmt3b* mirrored *Pou5f1*. This observation was further corroborated by independent long-read sequencing data (*66*), which showed that, while *Nacc1* and *Zfp281* expression peaked at the 2-Cell stage, *Dnmt3b* and *Sall4* peaked at the blastocyst stage (fig. S1E). These results suggest that *Sall4* and *Dnmt3b* may not be involved in inducing the TLC state in ESCs.

To determine whether *Nacc1*, *Sall4*, *Dnmt3b* and *Zfp281* are TLC regulators, we generated their individual knockouts (KOs) in ESCs using CRISPR-Cas9 (*67*) (see methods). To evaluate the effect of these KOs on TLCs, we utilized a well-established acetate treatment culture condition that induces TLCs in ESC cultures (*15, 21*) (see methods). We verified that acetate treatment efficiently induced the expression of TLC genes and increased the proportion of TLCs in comparison to pluripotency-promoting conditions (fig. S2A-C). To generate these KOs, we tested two guide RNAs (gRNAs) for *Nacc1* and three gRNAs for *Sall4*, *Dnmt3b* and *Zfp281*. As a positive control, we targeted *Duxf3*, a well-known TLC gene (*33, 34*), and as a negative control, we used an empty vector containing Cas9 without gRNA (henceforth referred to as the control). In agreement with its established role, *Duxf3* KO abolished the induction of TLCs with acetate treatment (fig. S2D). In contrast, *Sall4*, *Dnmt3b* and *Zfp281* KOs had minimal effects. However, we found that *Nacc1* KO drastically reduced the proportion of TLCs (fig. S2D), which prompted us to focus on its role in regulating totipotency.

To confirm *Nacc1* KO efficiency, we evaluated both mRNA and protein expression levels. gRNA2 turned out to be the best gRNA as its use resulted in near-complete KO (fig. S2E-F). To reduce heterogeneity, we derived clonal mutant cell lines, identifying five out of twelve clones with complete *Nacc1*-KO (fig. S2G). After confirming proper gene editing by Sanger sequencing, we selected clone2 (henceforth *Nacc1*^-/-^) for the rest of the study (fig. S2G-H). The proportion of TLCs was significantly reduced by *Nacc1*^-/-^ in acetate condition (Fig. 1C). Accordingly, *Nacc1*^-/-^ impaired the expression of multiple TLC genes in acetate and other TLC inducing conditions (fig. S2I) (*14, 15, 68–71*). In contrast, *Nacc1*^-/-^ effect on pluripotency genes was context-dependent: it reduced *Pou5f1*, *Sox2* and *Klf4* expression in SL and LB, as previously reported (*55, 72, 73*), but did not change in 2iL (fig. S2J). We next tested different acetate concentrations for the transition to the TLC state, and found that 40 mM was the optimal concentration (fig. S2K). Under these experimental conditions, the expression of TLC genes (*Zscan4, Duxf3*, *Zfp352*, *Tcstv1,* and MERVL) was induced in control cells at 40 mM acetate but was downregulated in *Nacc1*^-/-^ cells (Fig. 1D). On the other hand, overexpression of NACC1 was able to significantly increase the proportion of TLCs in lower concentrations of acetate, but had no effect at 40mM or in pluripotent conditions (fig. S2L-M).

### NACC1 is crucial for the expression of TLC and ZGA genes

To examine genome-wide changes in cell fate-specific gene expression, we performed RNA sequencing (RNA-seq) of control and *Nacc1*^-/-^ cells following acetate treatment and 2iL, TLC and pluripotency-promoting conditions, respectively (*15, 74*). Variance analysis showed a clear distinction between acetate- and 2iL-treated cells, and while *Nacc1*^-/-^ had a major effect on gene expression upon acetate treatment, it had a minor effect in 2iL (Figs. 2A and S3A). Differentially expressed genes (DEGs) revealed distinct condition-specific up- and down-regulation of transcripts. Compared to 2iL, acetate-treated cells were characterized by 7317 up-regulated genes, including the TLC genes *Zscan4b-f*, *Duxf3*, and *Zfp352*, and 4921 down-regulated genes, encompassing the pluripotency genes *Pou5f1*, *Lefty2*, *Nanog*, *Esrrb*, and *Klf2* (Fig. 2B). The distinct expression of TLC and pluripotency genes in acetate and 2iL conditions, respectively, further confirms their role in promoting specific cell fates (*15, 74*). While *Nacc1*^-/-^ in acetate-treated cells resulted in downregulation of 1989 genes, including *Nacc1* itself and TLC genes, and upregulation of 451 genes (Fig. 2C), gene expression in 2iL was barely affected (fig. S3B). Overall, a substantial proportion of genes upregulated in response to acetate (1760, i.e. 24%) were downregulated upon *Nacc1* knockout (Fig. 2D).

**Figure 2:**
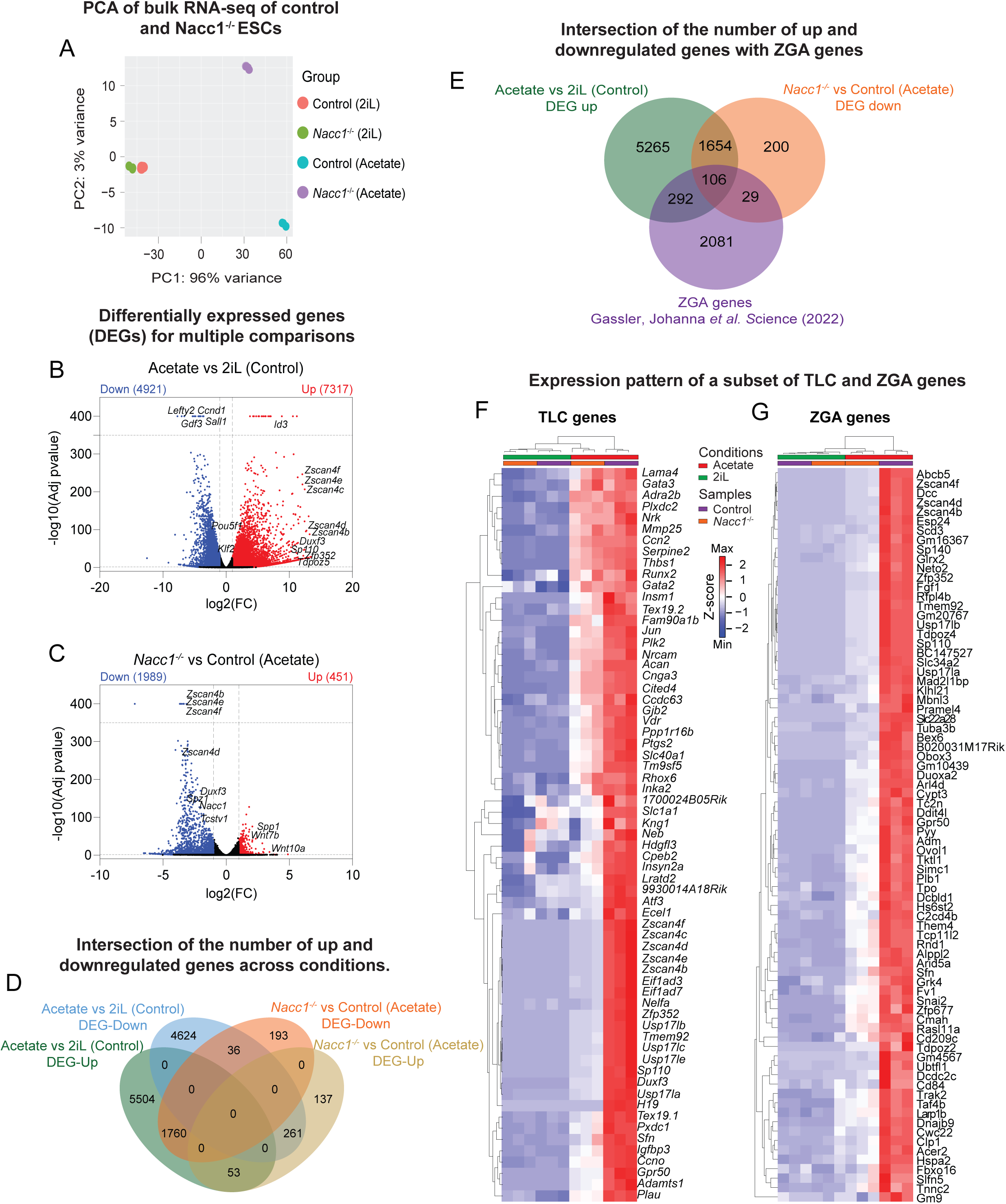
NACC1 is essential for the expression of TLC genes. **(A)** Principal component analysis of bulk RNA-seq of control and *Nacc1^-/-^* ESCs (n = 3) cultured in 2iL or acetate condition for 72h. **(B-C)** Volcano plots showing differentially expressed genes (DEGs) for acetate vs 2iL (control) (B), and *Nacc1^-/-^* vs control in acetate condition (C). Representative TLC and pluripotency genes are illustrated on the plots. Red dots indicate significantly upregulated DEGs, and blue dots indicate significantly downregulated DEGs. Grey line on X-axis indicates log2(FC)±1 and Y-axis indicates -log10(Adj p-value)=1.301 (lower limit) and 350 (upper limit). **(D)** Venn diagram showing the intersection of the number of up and downregulated genes in acetate vs 2iL (control), and *Nacc1^-/-^* vs control in acetate condition. **(E)** Venn diagram showing the intersection of upregulated genes in acetate vs 2iL (control), downregulated genes in *Nacc1^-/-^* vs control in acetate condition, and ZGA genes obtained from Gassler *et al.* (*75*) transcriptome datasets. **(F-G)** Hierarchically clustered heatmap showing the expression pattern of a subset of TLC genes (F) and 2-Cell blastomere genes (G) across control and *Nacc1^-/-^* ESCs cultured in 2iL or acetate. Columns represent individual samples. A subset of genes was obtained from Macfarlan *et al.* (*28*) and Gassler *et al.* (111).

To further evaluate whether ZGA and totipotency genes were NACC1 dependent, we compared our DEGs with curated TLC and ZGA genes (*28, 75*). Interestingly, expression of 135 ZGA genes turned out to be NACC1 dependent, out of which 106 genes were transcriptionally induced in response to acetate treatment (Fig. 2E). Furthermore, Z-score analysis revealed that the expression of TLC (*Duxf3*, *Zscan4b-f*, *Nelfa, Usp17la-c, Zfp352, Gata2….)* and ZGA signature genes (*Zscan4f/d/b*, *Esp24*, *Sp140*, *Zfp352, Tmem92, Snai2*….), was highly downregulated upon *Nacc1* knockout in acetate treated cells (Fig. 2F-G). As expected, these genes were not expressed in control and *Nacc1*^-/-^ cells in 2iL, where pluripotency genes remained largely unaffected, except for a few genes, such as *Inhbe*, *Pde1b,* and *Klf4* (fig. S3C).

Based on these results, we propose that NACC1 is crucial for the expression of the key totipotency and ZGA genes.

### NACC1 is critical for the expression of totipotency-associated retrotransposons

Given the association between totipotency and the activation of retrotransposons (*11, 37*), we explored the possible implication of NACC1 in modulating retrotransposon expression. This was achieved by performing a transposable element (TE)-focused analysis of our RNA-seq data in control and *Nacc1*^-/-^ cells cultured in acetate and 2iL. The analysis of differentially expressed transposable elements (DETEs) showed a strong induction of retrotransposon expression by acetate (relative to 2iL) in control cells, with 395 upregulated and only 10 downregulated TEs (Fig. 3A). *Nacc1* knockout resulted in 103 downregulated and only 2 upregulated retrotransposons, which were mainly unchanged in 2iL (Figs. 3B and S4A). Remarkably, the long-terminal repeat (LTR) class of retrotransposons (MERVL-int, MT2_Mm, MERVL_2A-int, etc.), which are specifically activated at the totipotency stage (2C blastomeres), were strongly upregulated in control cells and downregulated in *Nacc1*^-/-^ cells (Fig. 3A-B).

**Figure 3:**
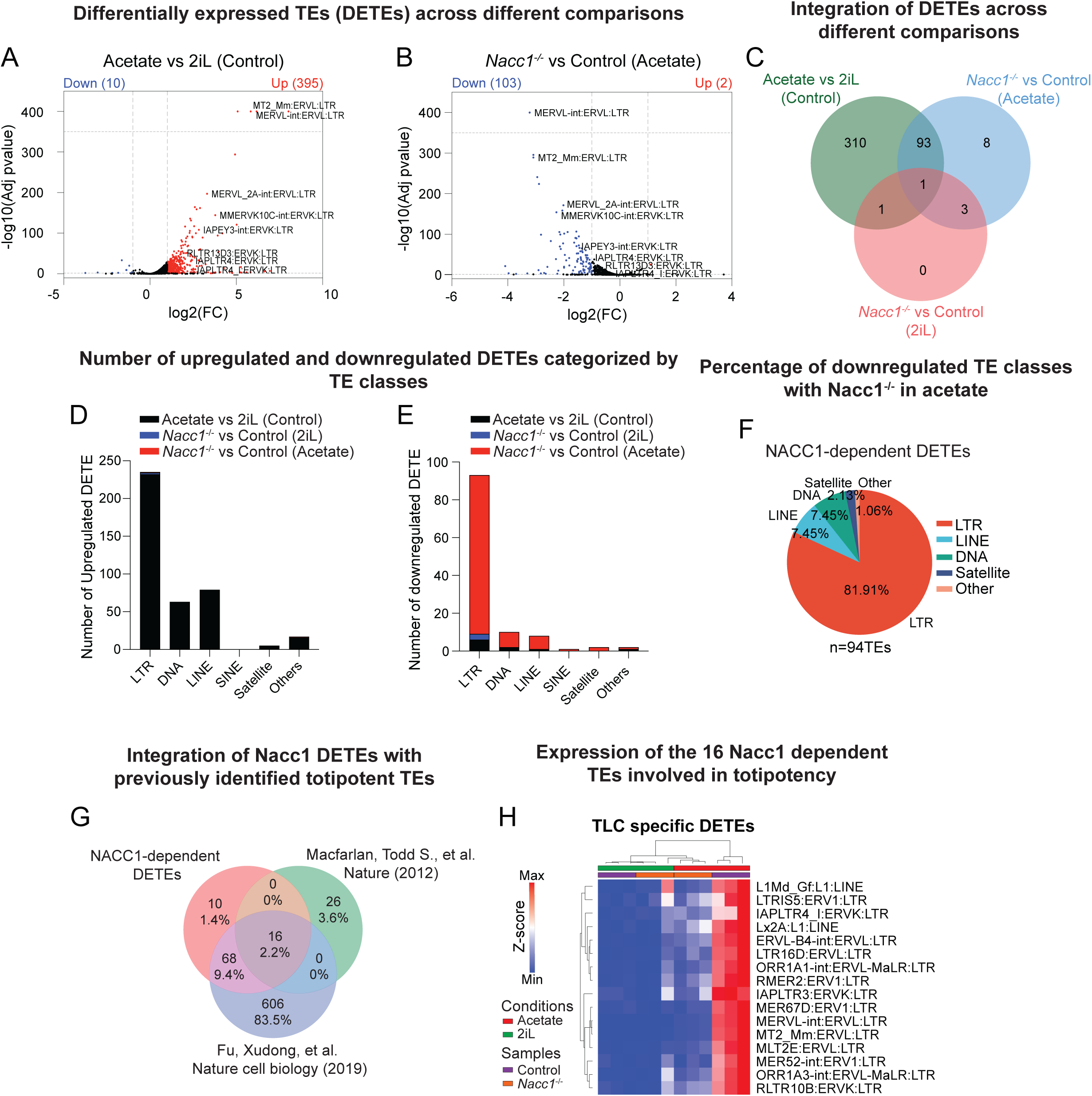
NACC1 is essential for the expression of retrotransposons. **(A-B)** Volcano plots showing differentially expressed transposable elements (DETEs) in acetate vs 2iL (control) (A), and *Nacc1^-/-^* vs control in acetate condition (B). Representative TLC associated retrotransposons are illustrated on the plot. Red dots indicate significantly upregulated TEs, and blue dots indicate significantly downregulated TEs. Grey line on X-axis indicates log2(FC)±1 and on Y-axis indicates -log10(Adj p-value)=1.301 (lower limit) and 350 (upper limit). **(C)** Venn diagram showing the intersection of the number of DETEs between acetate vs 2iL (control), *Nacc1^-/-^* vs control in 2iL, and *Nacc1^-/-^* vs control in acetate condition. **(D-E)** Bar plot showing the number of upregulated (D) and downregulated (E) DETEs categorized by TE subclasses in acetate vs 2iL (control), *Nacc1^-/-^*vs control in 2iL, and *Nacc1^-/-^* vs control in acetate condition. **(F)** Pie plot showing the percentage of downregulated TE subclasses with *Nacc1^-/-^* in acetate condition. **(G)** Venn diagram showing the comparison of upregulated TEs in 2C blastomeres (*27*), TLCs (*28*) and downregulated TEs in *Nacc1^-/-^* of acetate condition. **(H)** Hierarchical heatmap for the expression pattern of a subset of TEs enriched in TLCs (obtained from the intersection in (G)) across control and *Nacc1^-/-^*in 2iL or acetate condition comparisons. Columns represent individual samples.

Since *Nacc1* knockout had a major impact on TEs expression, we examined the overlap of DETEs across conditions and found 98 intersecting TEs; 93 for control and *Nacc1*^-/-^ in acetate treated cells, one for control and *Nacc1*^-/-^ in 2iL, one for all three groups and three for control and *Nacc1*^-/-^ in both acetate-treated and 2iL cells (Fig. 3C). To evaluate the nature of all DETEs, we annotated them by class and found that in both up and downregulated TEs, the majority belonged to LTR, followed by LINE and DNA transposons, while satellite and SINEs were nearly absent (Fig. 3D-E). Overall, a substantial proportion of TEs (94, corresponding to ∼24%) upregulated in acetate-treated cells were downregulated upon *Nacc1* knockout (fig. S4B). Among these 94 DETEs (∼82%) belonged to the LTR family, followed by LINE and DNA (∼7.5% each), satellite (∼2%), and others (∼1%) (Fig. 3F). To further identify the key NACC1-dependent TEs, we compared the 94 DETEs with two publicly available datasets of totipotency-related TEs (*11, 27*) and found 16 TEs in common (Fig. 3G). Among these 16 TEs, 14 are LTR elements (including MERVL-int and MT2_Mm), and two are LINE elements (Lx2A and L1Md_Gf). The Z-score expression profile of these 16 TEs further revealed that their expression was high in control cells and completely downregulated in *Nacc1*^-/-^ cells upon acetate treatment, while they were not expressed in either control or *Nacc1*^-/-^ cells in 2iL (Fig. 3H).

Taken together, these data reveal that NACC1 is critical for the expression of key totipotency-associated genes and retrotransposons.

### NACC1 binds the regulatory regions of both TLC genes and retrotransposons

NACC1 is known to directly bind DNA through its BEN domain, by recognizing the core consensus CATA/G sequence (*76*). It can bind to the promoter regions of its target genes to either maintain the pluripotency of ESCs or coordinate their differentiation (*55, 77*). Its binding to the regulatory regions of *Zeb1* and *E-cadherin* during somatic cell reprogramming promotes the establishment of the pluripotency state (*72*). Given this mechanism, we hypothesized that NACC1 could similarly interact with the regulatory regions of key totipotency genes and retrotransposons to induce their expression. To test this hypothesis, we performed chromatin immunoprecipitation followed by sequencing (ChIP-seq) in ESCs cultured in 2iL and acetate conditions. ChIP-seq peaks were categorized as acetate-enriched, 2iL-enriched, or common to both conditions, with a substantial number of peaks detected across all three groups (Fig. 4A). Examining the genomic distribution of these peaks revealed that for acetate-enriched regions, NACC1 binding was relatively enriched within 1 kb of the transcription start site (TSS) but reduced at 10–100 kb compared to 2iL-enriched peaks (Fig. 4B). Across all categories, NACC1 predominantly bound near promoter and distal intergenic regions (fig. S5A-C). *De novo* motif analysis using Homer confirmed that NACC1 binds coding gene targets via its core consensus sequence, regardless of the peak classification (Fig. 4C). Correlating with its requirement for expression of TLC genes, NACC1 binding to all TLC genes was relatively more enriched in acetate than in 2iL conditions (Fig. 4D).

**Figure 4:**
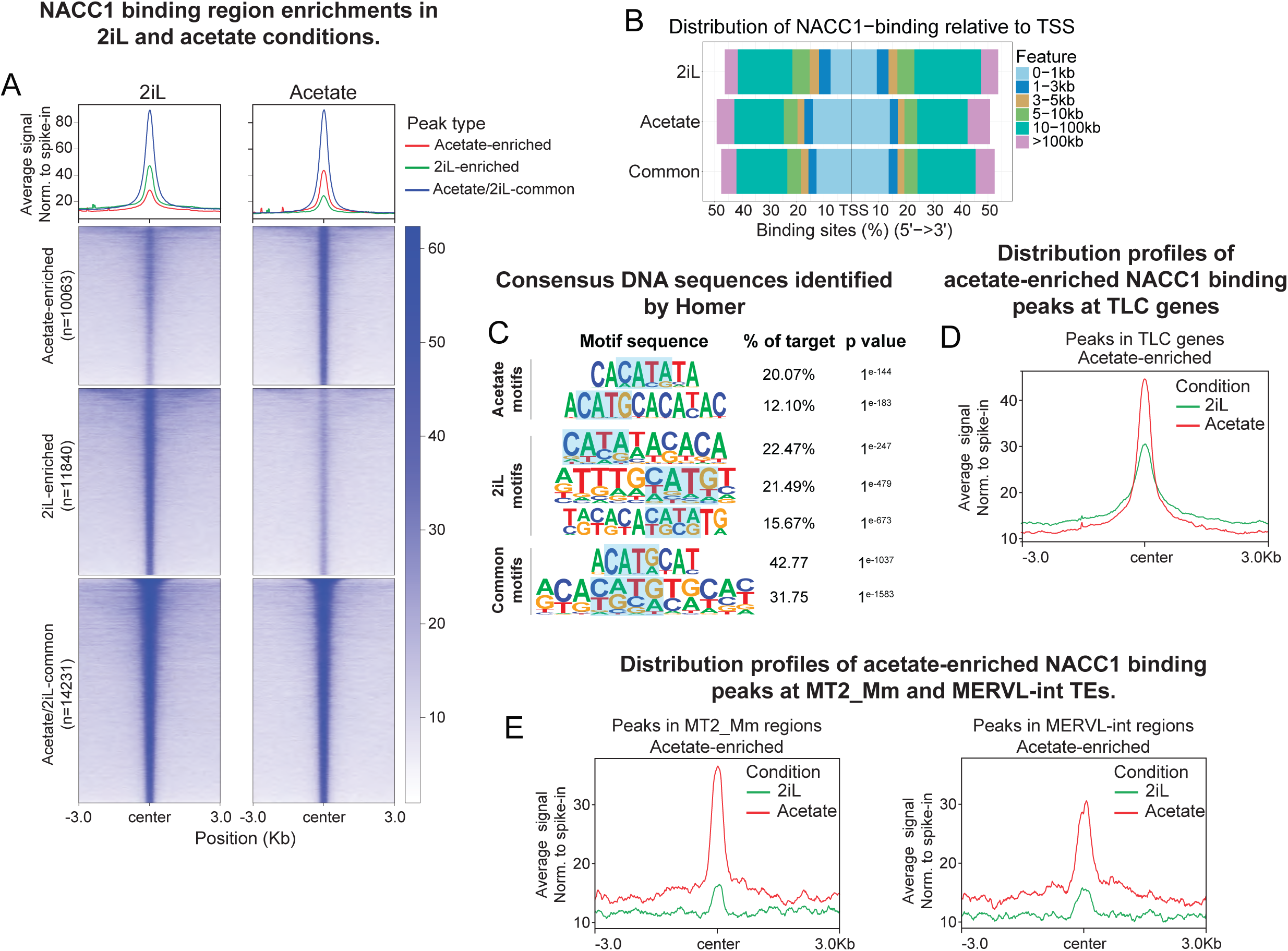
NACC1 directly binds regulatory regions of TLC genes and retrotransposons. **(A)** Line plots (above) and heatmaps (below) of NACC1 binding region enrichments in 2iL and acetate conditions. Average signals of merged replicates are shown. NACC1 peaks are classified by acetate-enriched (n=10063), 2iL-enriched (n=11840) and acetate/2iL-common (n=14231) regions. Signals were normalized using spike-in (Methods). **(B)** Distribution of the above classified NACC1-binding peaks according to the distance from the nearest transcription start site (TSS). **(C)** Consensus DNA sequences identified by Homer *de novo* motif analysis using NACC1 binding peaks. The corresponding percentage of target peaks and p-values are shown. Highlighted region represents the consensus core motif (CATC/G) of recognition for NACC1’s BEN domain. **(D-E)** Distribution profiles of acetate-enriched NACC1 binding peaks at TLC gene regions (D), and MT2_Mm and MERVL-int retrotransposons (E) in 2iL and acetate conditions.

Next, we determined whether NACC1 also binds directly to retrotransposons. We performed a TE-focused analysis of our ChIP-seq dataset. Motif analysis using Homer, MEME and FIMO confirmed that NACC1 also binds retrotransposon targets via its core consensus sequence (fig. S5D). The differential peak enrichment analysis based on spike-in showed that NACC1 binding at all TE classes is more enriched in acetate than 2iL (fig. S5E). We then examined whether NACC1 specifically binds to the 16 retrotransposon subclasses identified in our RNA-seq analysis as NACC1-dependent (Fig. 3H). We detected NACC1 binding in 10 of these 16 retrotransposons. For instance, we observed significant enrichment of NACC1 binding at MT2_Mm and MERVL-int regions specifically in acetate-treated cells (Figs. 4E and S5F).

Overall, these data indicate that NACC1 directly binds regulatory regions of both totipotency genes and retrotransposons.

### NACC1 promotes chromatin accessibility of its targets

To determine whether NACC1 is required for the chromatin accessibility of its targets, we performed assays for Transposase-Accessible Chromatin using sequencing (ATAC-seq) on control and *Nacc1*^-/-^ cells cultured in 2iL and acetate conditions. To gain insights into differences in accessibility across conditions and *Nacc1* dependency, we analyzed ATAC-seq peaks for differentially accessible regions (DARs) using DESeq2 (see methods). Based on the relative ATAC-seq peak signal strength, the DAR analysis between samples reveals differential chromatin accessibility, with high signal representing accessible (or open) region and low or no signal representing non-accessible (or closed) region. Chromatin accessibility differed strikingly between the control and *Nacc1* knockout cells in the acetate condition, with a drastic reduction in the signal of accessible regions in the latter (Fig. 5A). In comparison to control cells in the acetate condition, these regions were relatively non-accessible in control and *Nacc1* knockout cells in the 2iL condition. In analyzing the TLC gene-specific regions, we found that their accessibility was prominent in acetate compared to the 2iL condition (Fig. 5B). In acetate condition, *Nacc1* knockout resulted in a drastic reduction in the accessibility of TLC gene regions. On the other hand, *Nacc1* knockout had least impact on the accessibility of TLC gene regions in the 2iL condition (Fig. 5C).

**Figure 5:**
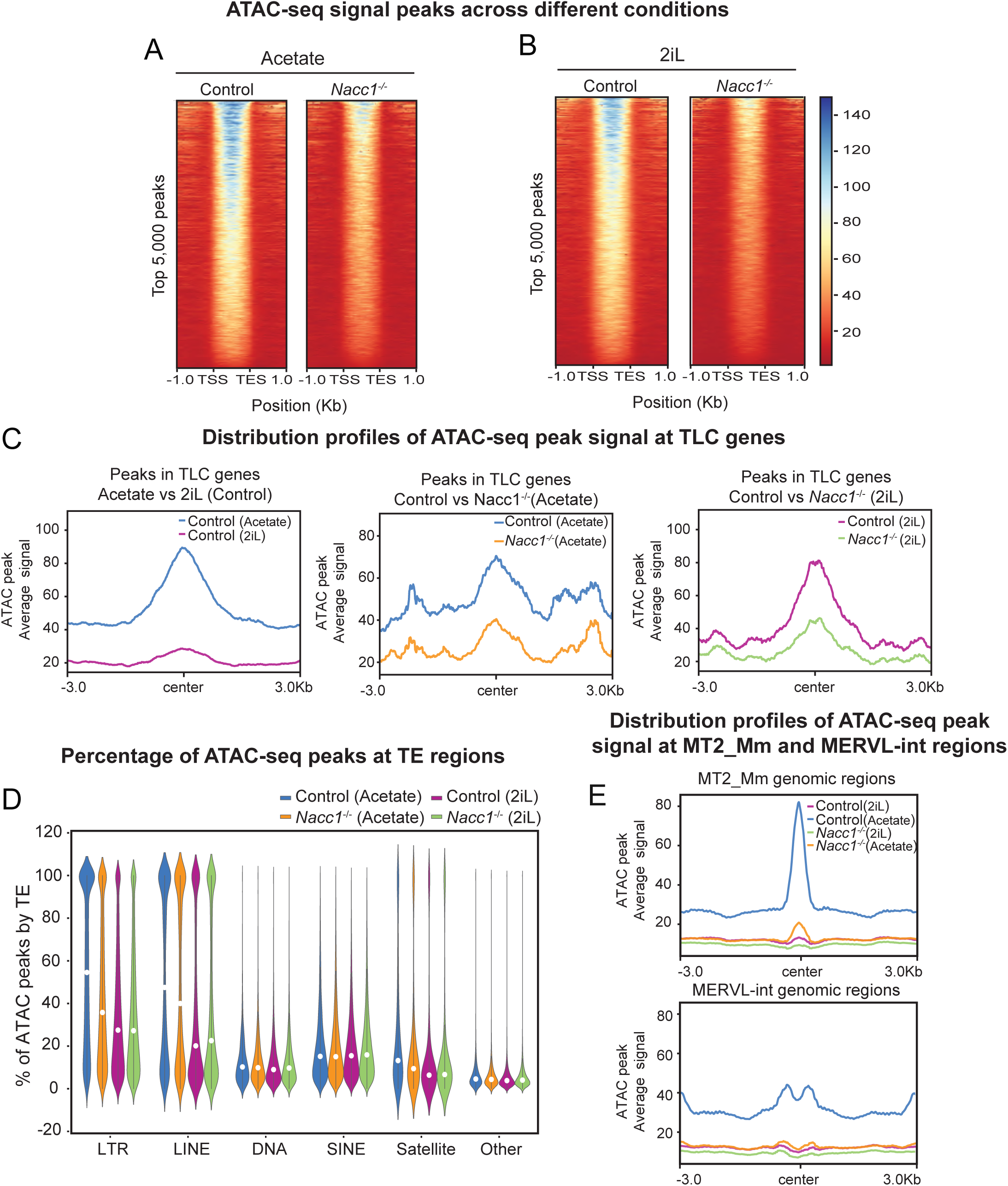
NACC1 promotes chromatin accessibility of TLC genes and retrotransposons. **(A-B)** Heatmaps showing the average ATAC-seq signals of top 5000 open DARs per condition in control vs *Nacc1^-/-^* in acetate (A), and control vs *Nacc1^-/-^* in 2iL (B). **(C)** Distribution profiles of ATAC-seq peaks at TLC genes in acetate vs 2iL (control), *Nacc1^-/-^* vs control in acetate, and *Nacc1^-/-^* vs control in 2iL condition. **(D)** Violin plot showing the percentage of ATAC-seq peaks categorized by TE-subclass in control and *Nacc1^-/-^* samples analyzed in 2iL or acetate conditions. **(E)** Distribution profiles of ATAC-seq peaks at all genomic regions of MT2_Mm and MERVL-int retrotransposons in control and *Nacc1^-/-^*in acetate and 2iL conditions.

Next, we analyzed chromatin accessibility at retrotransposon-associated regions. The analysis revealed that compared to control in acetate-treated cells, *Nacc1* knockout led to a reduction of ATAC peaks/signal, specifically in LTR, LINE, and satellite TEs, whereas no significant effect was observed across these retrotransposon classes in 2iL (Fig. 5D). The accessibility at specific retrotransposons, such as MT2_Mm and MERVL-int, was increased in control acetate-treated cells, and this effect was abolished upon *Nacc1* knockout (Fig. 5E). In contrast, the MT2_Mm and MERVL-int regions remained non-accessible in 2iL cells, with no significant effect associated with *Nacc1* knockout (Fig. 5E).

These ATAC-seq data suggest that NACC1 facilitates chromatin accessibility of both totipotency genes and retrotransposons.

### NACC1 targets both TLC genes and retrotransposons to regulate their expression

To further understand the impact of NACC1-dependent chromatin accessibility on gene expression, we intersected the promoter-specific accessibility and gene expression observed in our ATAC-seq (log2FC±0.5, FDR < 0.05) and RNA-seq (log2FC±1, FDR < 0.05) datasets, respectively. As anticipated, gains of chromatin accessibility overlapped with upregulated genes, while losses overlapped with downregulated genes (Figs. 6A-B and S6A). In control cells, accessible genes with increased expression included TLC genes such as *Duxf3, Zscan4f*, *Usp17le*, *Zfp352, Tcstv3, Rarg* and *Hand1*, whereas non-accessible genes with decreased expression included pluripotency genes such as *Pou5f1, Zfp42*, *Lefty2*, *Dnmt1* and *Lefty1* (Fig. 6A). Accordingly, *Nacc1* knockout in acetate-treated cells led to loss of chromatin accessibility and downregulation of TLC genes (e.g. *Duxf3*, *P4ha2*, and *Fkbp6),* and lesser effect was observed in 2iL (Figs. 6B and S6B). Consistently, IGV tracks at representative loci showed NACC1 binding predominantly at TLC genes (e.g., *Duxf3*, *Usp17le*, *Gm4340,* and *Zscan4d*), along with their enhanced chromatin accessibility and increased expression. This effect was abolished in *Nacc1*^-/-^ cells (Fig. 6C). Also, in agreement with its limited impact in 2iL, *Nacc1* knockout did not significantly alter chromatin accessibility or expression of TLC or pluripotency genes in 2iL (Figs. 6C and S6C). Interestingly, while NACC1 binding was relatively enriched at pluripotency genes (e.g., *Lefty2, Nanog and Pou5f1*) in 2iL, their chromatin accessibility and expression remained NACC1-independent (fig. S6C).

**Figure 6.**
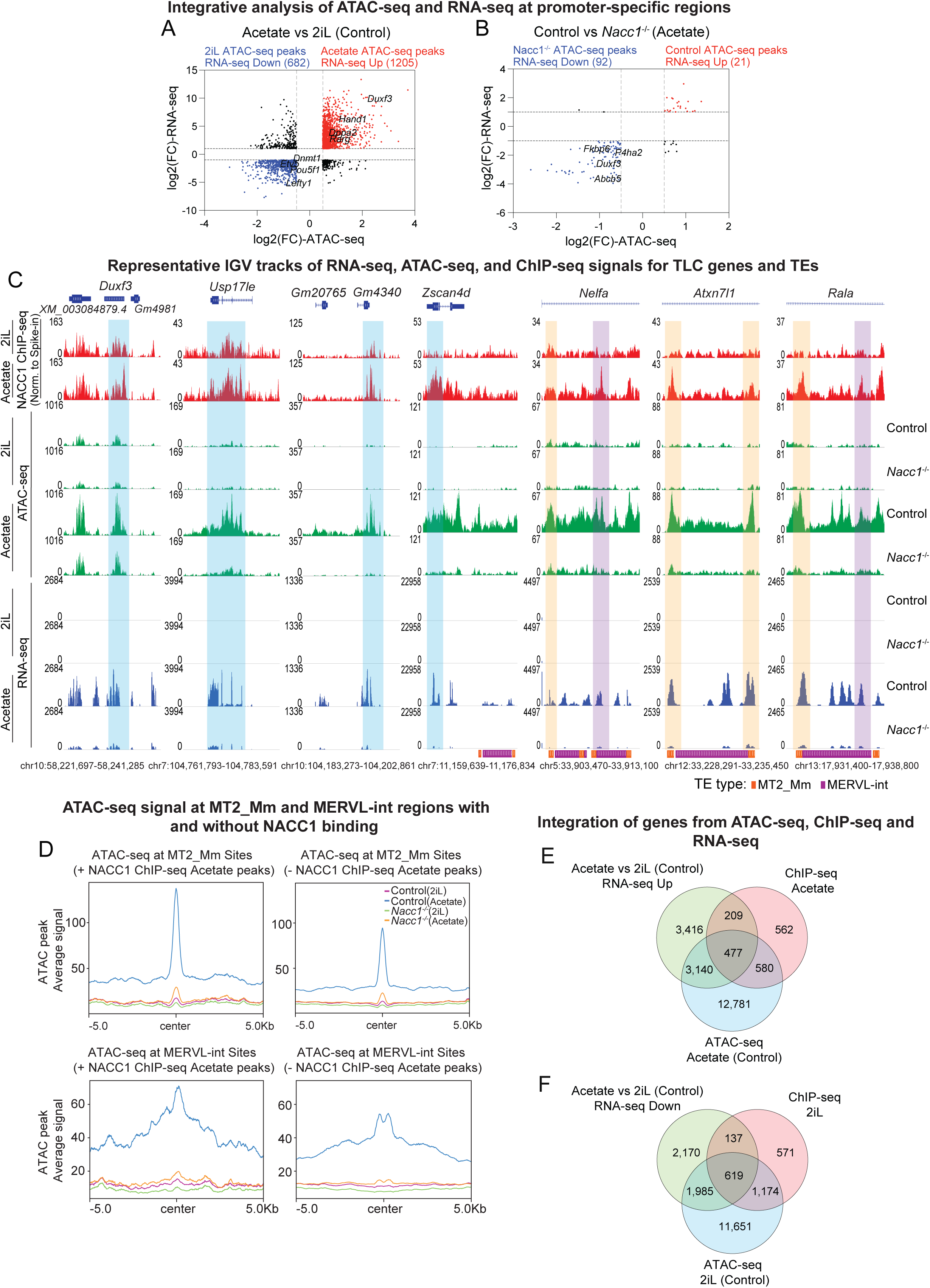
NACC1 targets both TLC genes and retrotransposons to regulate their expression. **(A-B)** Integrative analysis of DARs and DEGs at promoter-specific regions in acetate vs 2iL (control) (A) and *Nacc1^-/-^* vs control in acetate condition (B). Red dots indicate significantly upregulated genes with open accessible chromatin regions, and blue dots indicate significantly downregulated genes with closed non-accessible chromatin regions. Grey lines indicate log2(FC)±0.5 on X-axis and log2(FC)±1 on Y-axis. Representative TLC and pluripotency genes are illustrated on the plots. **(C)** Representative IGV snapshots of RNA-seq, ATAC-seq and ChIP-seq signals for TLC genes (*Duxf3, Usp17le*, *Gm4340* and *Zscan4d*) and for MT2_Mm (orange) and MERVL-int (purple) retrotransposons proximal to TLC genes (*Zscan4d, Nelfa, Atxn7l1, Rala*). Boxes indicate changes in the peaks for TLC genes (blue), MT2_Mm (orange) and MERVL-int (purple). **(D)** Distribution profiles of chromatin accessibility peaks at MT2_Mm and MERVL-int regions with and without NACC1 enrichment using acetate-enriched peaks from ChIP-seq. **(E-F)** Venn diagrams showing the gene integration of ATAC-seq, ChIP-seq and RNA-seq datasets from acetate (E) or 2iL (F) control samples.

Analyzing the intersection of ChIP-seq and ATAC-seq peaks for MT2_Mm and MERVL-int revealed that overlapping peaks were observed in acetate but not in 2iL conditions (fig. S6D-E). The marked enrichment of ATAC-seq peak signal at NACC1-bound regions in acetate for MT2_Mm and MERVL-int was lost upon *Nacc1* knockout (Fig. 6D). Furthermore, chromatin accessibility was lost upon *Nacc1* knockout at MT2_Mm and MERVL-int loci which are proximally located (proximal) to TLC genes such as *Zscan4d*, *Nelfa, Atxn7l1* and *Rala* (Fig. 6C). This loss of accessibility strongly correlated with high NACC1 binding at the MT2_Mm and MERVL-int loci in control cells, and loss of their and TLC genes expression in acetate-treated *Nacc1* knockout cells (Fig. 6C). Integrating ATAC-seq, RNA-seq, and ChIP-seq data revealed that 477 and 619 genes were uniquely regulated by NACC1 in acetate and 2iL conditions, respectively (Fig. 6E-F).

Overall, these results demonstrate that NACC1 induces the expression of both TLC genes and retrotransposons by directly binding to their regulatory regions and promoting their chromatin accessibility.

### NACC1-regulated retrotransposons modulate the expression of proximal genes

Retrotransposons can regulate the expression of proximal genes in the genome (*78–81*). To test whether NACC1-regulated retrotransposons modulate the expression of proximal genes, including TLC genes, we focused on the 16 retrotransposons strongly regulated by NACC1 (Fig. 3H). We performed proximal gene regulation analysis by overlapping differentially expressed genes from our transcriptome data with the repeat annotation available on RepeatMasker (see Methods) (*82*). We found that three of the 16 retrotransposons (MERVL-int, MT2_Mm, and ORR1A3-int) are predicted to modulate proximal gene expression in the control acetate condition (Figs. 7A and S7A). A similar pattern of proximal regulation was observed among NACC1-dependent genes in the acetate condition (Figs. 7B and S7B). All three retrotransposons have been linked to the TLC state (*28*). Interestingly, they appeared to control the expression of ∼7.5 to 12.5% of upregulated genes in control samples and ∼4 to 10% of NACC1-dependent genes in acetate (Fig. 7A-B). The predominant proportion of genes was regulated by MT2_Mm, followed by MERVL-int and ORR1A3-int. Consistent with the absence of NACC1-mediated regulation of retrotransposons in 2iL (fig. S4A), proximal regulation was not observed for NACC1-dependent genes in 2iL (fig. S7C-D).

**Figure 7:**
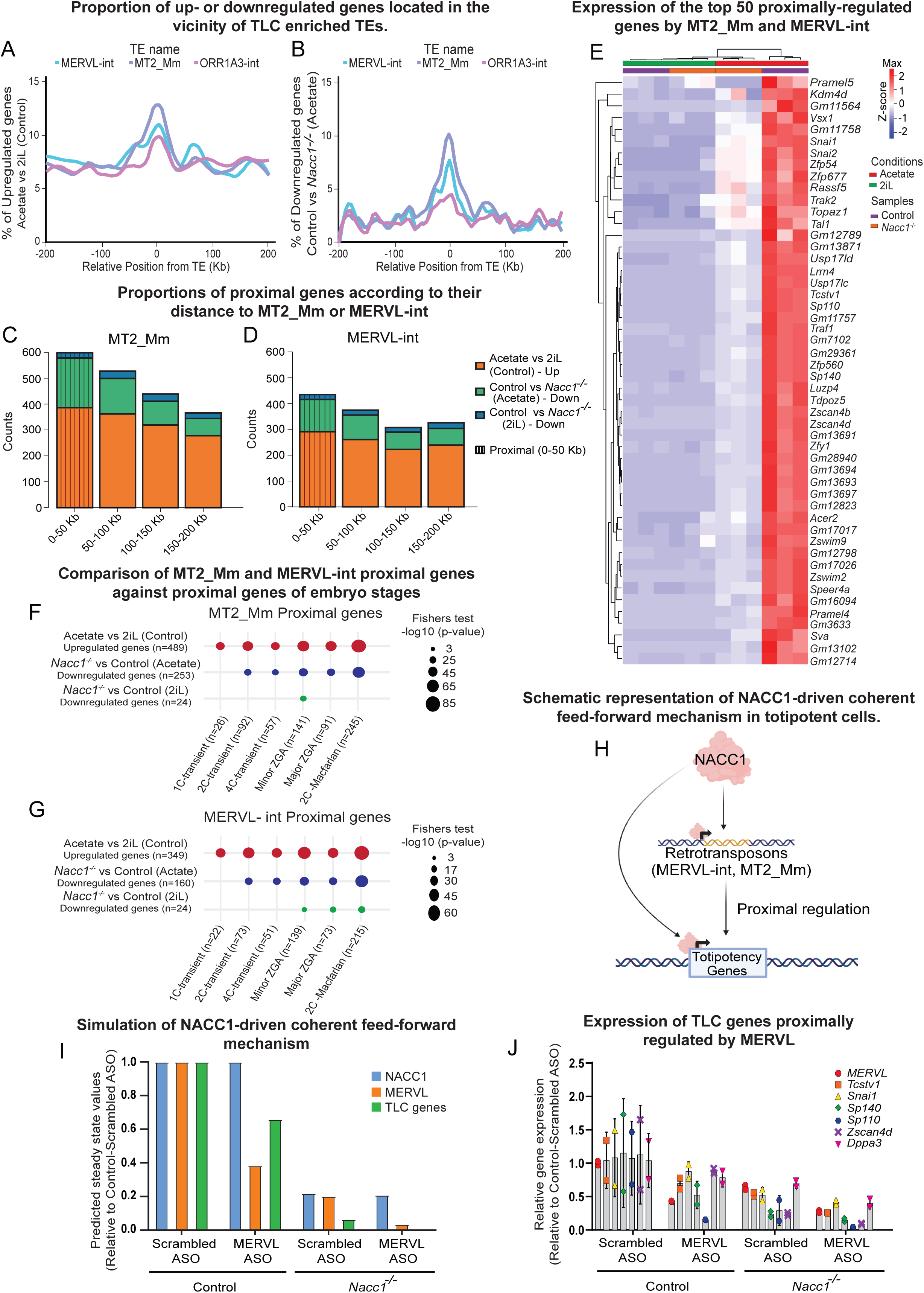
Retrotransposon-mediated regulation of proximal TLC genes is NACC1-dependent. **(A-B)** Proportion of up- or downregulated genes located in the vicinity of specific TLC enriched TEs. Lines represent the percentage trend of upregulated genes (acetate vs 2iL (control)) (A), and downregulated genes (*Nacc1^-/-^* vs control in acetate) (B) among all overlapping genes up to a distance of 200 kb surrounding each retrotransposon type. Blue, MERVL-int; purple, MT2_Mm; pink, ORR1A3-int (Methods). **(C-D)** Proportions of up- and downregulated proximal genes according to their distance to MT2_Mm (C) or MERVL-int (D) retrotransposons. Very proximal (0-10 Kb) and proximal regulation (10-50 Kb) proportions are specified by diagonal and straight lines, respectively, within a sample group. **(E)** Heatmap showing the expression pattern of the top 50 proximally regulated TLC genes across control and *Nacc1^-/-^* samples in 2iL or acetate conditions. Columns represent individual samples. **(F-G)** Bubble plot comparing the proximal genes regulated by MERVL-int (F) or MT2_Mm (G) within upregulated genes of acetate vs 2iL (control), and downregulated genes of *Nacc1^-/-^* vs control in acetate and 2iL conditions, against proximal genes regulated by the respective retrotransposon in Macfarlan *et al.* (*28*) and DBTMEEv2 (*75*) transcriptome datasets. The size of the dot depicts the –log10[p-value] derived from Fisher’s t-test. **(H)** Schematic representation of NACC1-driven coherent feed-forward mechanism in totipotent cells. **(I)** Predicted steady-state concentrations of NACC1 (blue), MERVL (orange), and TLC genes (green) in wild type or knockout/knockdown conditions using an ODE simulation. **(J)** RT-qPCR analysis for the expression of proximally-regulated TLC genes by MERVL-int in control and *Nacc1^-/-^* cells treated with scrambled or MERVL ASOs in acetate condition for 48h. All samples were normalized to the scrambled ASO-treated control cells.

For both MERVL-int and MT2_Mm, the number of proximally regulated genes decreased as the distance from the retrotransposon increased, regardless of the condition, with the highest number of genes found up to 50 kb away (Fig. 7C-D). Although defining the precise distance at which proximal regulation occurs remains challenging, previous studies have documented retrotransposons influencing gene expression up to a couple of hundred kb away, through chromatin looping (*78, 83*). Following this observation, we categorized genes within 0-10 kb as very proximal, as this range captures direct interactions in *cis*, and genes within 10-50 kb as proximal, as they could still be proximally regulated through potential enhancer-mediated or chromatin-looping interactions (*78, 83*).

Among the NACC1-dependent proximally regulated genes in acetate-treated cells, a significant number of genes were regulated by MT2_Mm and MERVL-int (fig. S8E-F). Genes commonly regulated by both MERVL-int and MT2_Mm were maximally expressed in control cells but were drastically downregulated in *Nacc1*^-/-^ cells with acetate (e.g., *Zscan4b*, *Zscan4d*, *Tdpoz5, Tcstv1, Sp110, Sp140, Zfp54, and Gm12789*) (Figs. 7E and S7G). To test whether retrotransposon-regulated proximal genes are expressed in the earliest stages of preimplantation embryo development (*28, 81*), we compared proximally regulated genes of MT2_Mm and MERVL-int identified in our dataset with proximally regulated genes of 2C genes (as defined in Macfarlan et al.(*28*)), and 1C-4C and ZGA gene sets from DBTMEE v2 (*75*). Significant enrichment was observed for proximal genes upregulated by acetate with all gene sets (Fig. 7F-G). Similarly, significant enrichment was identified for NACC1-dependent proximal genes in acetate with 2C, 4C, and ZGA genes. Such enrichment was not observed with 1C genes. In comparison, NACC1-dependent proximal genes in 2iL had minimal to no enrichment (Fig. 7F-G).

### Regulation of TLC genes by NACC1-driven feed-forward mechanism

These results imply that NACC1 regulates TLC genes through a coherent feed-forward loop, acting both directly or indirectly through transposable elements such as MERVL or MT2_Mm (Fig. 7H). To predict the dynamic behaviour of this network, we implemented an ordinary differential equation (ODE) model (see Methods) and simulated the temporal and steady-state expression of each component under normal or perturbed conditions (Fig. 7I). In the wild-type simulation, all components reached steady state in a secuential manner: NACC1 accumulated first at a relatively low level, which lead to the activation and stabilization of MERVL, ultimately driving the induction of TLC genes (fig. S7H). Interestingly, perturbation of either NACC1 or MERVL, using experimentally observed KO or KD values, delayed TLC gene induction and reduced the steady-state levels of all components, with the strongest effect observed upon NACC1 removal. Furthermore, reduction of both NACC1 and MERVL abolished TLC gene induction entirely (Figs. 7I and S7H).

To experimentally validate the proximal regulation of key TLC genes and feed-forward loop regulation, we employed a previously published approach of anti-sense oligo (ASO)-mediated knockdown (KD) of MERVL-int in control and *Nacc1*^-/-^ cells under acetate conditions (*39*). Notably, relative to scrambled ASO, MERVL KD resulted in partial downregulation of most of the proximal TLC genes in control samples as inferred previously by our ODE simulation (Fig. 7J). While the expression of MERVL and most TLC genes was also partially downregulated, *Sp140, Sp110* and *Zscan4d* were nearly completely downregulated in *Nacc1*^-/-^ cells. However, *Nacc1* knockout, in combination with MERVL KD, resulted in near-complete downregulation of all proximal TLC genes as expected (Fig. 7J). In addition to confirming the proximal gene regulation by MERVL, these results revealed both individual and parallel requirements of NACC1 and MERVL in controlling the expression of TLC genes.

Together, these data suggest that NACC1 controls the expression of TLC genes via two parallel pathways of a feed-forward mechanism: 1) by direct induction of TLC gene expression and 2) indirectly by amplifying their expression through retrotransposon-mediated proximal regulation. This NACC1-dependent mechanism appears crucial for establishing TLCs, with transcriptomes resembling those of the 2C stage of preimplantation embryos.

### NACC1 is required for early embryonic development

Given that NACC1 is required for the TLC state in ESCs and that its feed-forward regulatory targets are involved in early developmental processes, we reasoned that NACC1 might be crucial for early embryonic development. To first test this *in vitro*, we either treated sorted TLCs with or without NACC1 inhibitor (NIC3) or used sorted TLCs generated from *Nacc1*^-/-^ and assessed their ability to develop into blastoids (blastocyst-like structures) (*12*) (Fig. 8A). After 7 days in culture, inhibition of NACC1 significantly reduced blastoid formation at comparable levels as *Nacc1*^-/-^ (Fig. 8B). This result suggested that NACC1 is necessary for the developmental progression of TLCs into blastocyst-like structures *in vitro*.

**Figure 8:**
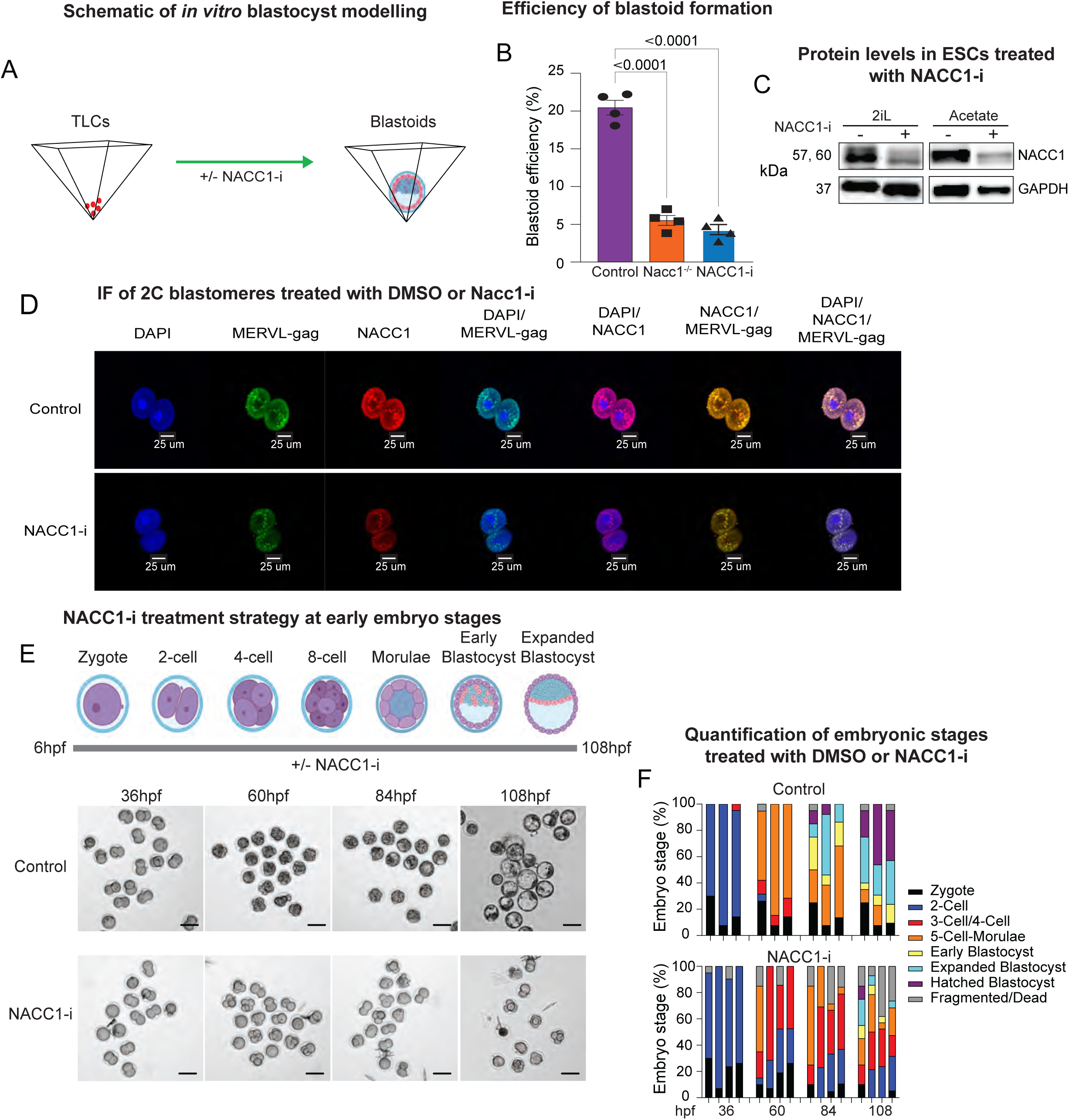
NACC1 is required for early stages of preimplantation embryo development. **(A)** Schematic of *in vitro* blastocyst modelling (blastoid formation) using WT or *Nacc1^-/-^*TLCs. Sorted TLCs were treated with either DMSO (control) or *Nacc1* inhibitor (NIC3, 20 μM). **(B)** Quantification of the efficiency of blastoid formation from *Nacc1^-/-^* TLCs or TLCs treated with either DMSO (control) or NIC3 (20 μM). Graph represents mean ± SEM, n = 4. *P*-value was determined by one-way ANOVA. **(C)** Western blot for NACC1 and GAPDH in ESCs treated with NACC1 inhibitor NIC3 in 2iL and acetate conditions. **(D)** Fluorescence images of DAPI, MERVL-gag, and NACC1 immunostained 2C blastomeres treated with DMSO (control) or NIC3 (20 μM) for 24h. Scale bars, 25μm. **(E)** Schematic for NIC3 treatment strategy at early embryo stages and stereomicroscopic images of the embryonic stages observed at the indicated time points under DMSO (control) and NIC3 (20μM) conditions. Scale bars, 100μm. **(F)** Quantification of the embryonic development stages from images in D. Sample sizes are as follows: mock/control (upper panel), n = 20, 18 and 21 embryos; NIC3 (20μM) (bottom panel), n = 20, 14, 21 and 19 embryos.

To further confirm NACC1 requirement for early embryonic developmental progression, we performed *ex vivo* embryo culturing from zygote-to-blastocyst stages in the presence or absence of NIC3. For this, we first confirmed that NIC3 treatment decreases NACC1 levels in 2C blastomeres (Fig. 8C). We treated the zygotes with either DMSO (control) or NIC3 (20 μM), fixed them at 2C blastomeres and immunostained for NACC1 and MERVL-gag. We observed relatively high levels of NACC1 in control and its expression level was drastically reduced in NIC3-treated blastomeres (Fig. 8C). Correlating with the change in NACC1 level, we observed a decrease in MERVL-gag levels from control to NIC3-treated blastomeres. Next, to assess the impact of NACC1 inhibition on developmental progression, we treated zygotes with either DMSO or NIC3 and allowed them to develop for 4.5 days, following which we quantified various phenotypes (Fig. 8D). Control blastomeres exhibited normal, timely developmental progression to form blastocysts (Fig. 8D-E). In sharp contrast, the majority of the NIC3-treated blastomeres displayed developmental arrest at 3-Cell to morula stages, with very few progressing to the blastocyst stage (Fig. 8D-E).

Together, these data demonstrate that NACC1 is crucial for the establishment of the totipotency stage in a mammalian embryo and its subsequent developmental progression to form a proper blastocyst.

## DISCUSSION

In this study, we identified NACC1 as a key regulator of TLCs by demonstrating its requirement to activate the expression of both totipotency genes and retrotransposons. Our data show that NACC1 directly binds to the regulatory regions of totipotency genes and retrotransposons, promoting chromatin accessibility and inducing their expression. Moreover, we identify a coherent feed-forward mechanism through which NACC1 promotes TLCs by directly activating totipotency genes and amplifying their expression via retrotransposon activation.

Our work expands the repertoire of NACC1 functions by demonstrating its critical role in totipotency, in addition to its well-established function in pluripotency (*55, 72, 73*). NACC1 is a component of the pluripotency TF network, known to promote pluripotent stem cell maintenance and required for somatic cell reprogramming to iPSCs (*55, 72, 73, 84, 85*). In serum with LIF culture condition, NACC1 promotes pluripotency by inducing the expression of *Nanog*, *Oct4,* and *Sox2,* while repressing *Tcf3* (*55*). It also plays context-dependent roles in ESC differentiation (*55*). In disease contexts, NACC1 contributes to tumour proliferation, promotes epithelial-to-mesenchymal transition in ovarian and pancreatic cancers (*86–90*), and is implicated in autoimmunity and neurodegenerative diseases such as Parkinson’s and Amyotrophic Lateral Sclerosis (*91–93*). The diverse functions of NACC1 may arise from its BEN (BANP, E5R and NACC1) and BTB/POZ (bric-a-brac/poxvirus and zinc finger) domains, which enable it to bind DNA and interact with other proteins to potentially form context-specific gene regulatory complexes (*59, 76, 94–97*). Our discovery of NACC1’s role in totipotency deepens our understanding of early embryonic cell fate specification, which should help in better understanding its role in disease conditions, where similar feed-forward mechanisms could play a central role in regulating cell plasticity and differentiation. Altogether, *Nacc1* appears to play a multifaceted role in regulating cell fates during early embryogenesis and in the context of diseases.

Transposable elements have significantly influenced the evolution of the mammalian genome through genetic innovation and regulation of gene expression (*38, 78, 98*). Nearly 40 to 45% of the mammalian genome corresponds to transposable elements, of which the majority are retrotransposons or endogenous retroviruses (*99, 100*). Recently, retrotransposons have been documented to regulate preimplantation embryo development (*28, 37, 39*). In particular, LTR retrotransposons such as MERVL and MT2_Mm are activated at the 2C stage of totipotency, and inhibiting their expression causes embryonic lethality and developmental arrest beyond the morula stage (*28, 37, 39*). Despite their importance, little is understood about how the expression of retrotransposons is controlled. Our data reveal the critical role of NACC1 in inducing the expression of different classes of retrotransposons, predominantly LTRs (∼82%, e.g., MERVL, MT2_Mm, and ORR1A3). NACC1 induced retrotransposon expression by directly binding to their DNA regulatory regions and promoting their chromatin accessibility. Among the LTRs, NACC1 prominently activated MT2_Mm elements, which can further act as promoters for MERVL and totipotency genes (*37*), thus positioning NACC1 as a pivotal regulator of totipotency. In agreement, we found that low NACC1 levels hinder the normal establishment of totipotency stage blastomeres and their subsequent developmental progression to blastocyst embryos. So far, few TFs have been reported to activate retrotransposon expression (*40, 41*), and, as reported here, the identification of NACC1 as one of them should be valuable for future studies aimed at better understanding retrotransposon regulation in development and disease.

Feed-forward motifs are recurrent network motifs found across diverse organisms, biological systems, and computation (*48, 49, 101–103*). With a defined information-processing function, feed-forward network motifs often act as core regulatory circuits or building blocks of complex networks (*48, 101, 102*). In this study, we describe the NACC1-driven coherent feed-forward mechanism as a pivotal regulatory process in totipotent cells, the initiators of mammalian embryo development. This mechanism involves NACC1 directly activating totipotency genes and amplifying their expression through retrotransposon activation. Although such network motifs have been documented for stem cell maintenance and differentiation (e.g., with OCT4, NANOG, and ESRRB in ESCs, and GATA1/2 in hematopoiesis) (*53, 54, 56, 57, 104*) and disease contexts (e.g., with NF-kB in adult T-cell leukemia, and MYC and PKC in cancer) (*105–107*), our report is the first instance of a feed-forward mechanism driving totipotency. NACC1 has been observed in feed-forward motifs with TCF3, OCT4, and SOX2 to regulate ESC differentiation into mesendoderm (*55*), further emphasizing its ability to orchestrate early embryogenesis through conserved network motifs. Our data also show that NACC1 activates DUX, ZSCAN4, and NELFA, well-established positive regulators of totipotency (*33, 34, 36, 108*), thereby situating them downstream of the NACC1 network. As feed-forward motifs can be nested or interlocked to regulate multiple cell fates during lineage specification (*48, 52, 109*), we speculate that NACC1 is likely to form such nested networks in cooperation with other regulators of totipotency, such as DUX, NELFA and ZSCAN4, to add robustness in establishing totipotency and pluripotency during preimplantation development.

Although NACC1 knockout in mice does not cause complete lethality, the survival rate of embryos is low and manifests developmental defects, highlighting NACC1’s role in early embryogenesis (*110*). Our data confirm that NACC1 is required for blastoid formation and the developmental progression of blastomeres from totipotency to the blastocyst stage. Its involvement in both totipotency and pluripotency suggests that NACC1 plays a critical role in regulating key stages of preimplantation embryogenesis. Gene regulatory mechanisms controlling the establishment of totipotency and its transition to pluripotency are poorly understood. Recently, an orphan nuclear receptor, NR5A2, was shown to regulate ZGA and is required for proper blastocyst development (*111*). Similar to NR5A2, NACC1 promotes chromatin accessibility and directly binds to gene-regulatory regions to induce the expression of ZGA genes. Moreover, NACC1 also significantly activates the expression of both totipotency genes and LTR retrotransposons associated with totipotency. Due to their similar and disparate roles, it is conceivable that NACC1 may function cooperatively with NR5A2, totipotency (e.g. DUX and NELFA) and pluripotency (e.g. OCT4, SOX2 and NANOG) factors to orchestrate mammalian preimplantation development. Besides the ZGA genes and SINE B1 TE, it is not yet clear whether NR5A2 similarly activates the specific bona fide totipotency-associated genes and retrotransposons.

In conclusion, this study uncovers a pivotal feed-forward gene regulatory mechanism governed by NACC1, a TF previously known for its role in pluripotency, which we show here is also crucial for totipotency. Our data demonstrate how a TF (e.g. NACC1) can simultaneously activate parallel pathways (e.g. totipotency genes and retrotransposons) to ensure robust induction of cell fate-specific gene expression programs. Investigating additional mechanisms implicated in the cooperative function of NACC1 and its TF partners will be important to deepen our understanding of the molecular mechanisms underpinning totipotency and may potentially lead to the utilization of TLCs for disease modelling and regenerative medicine.

## Acknowledgements

We are grateful to Marie Kmita, Nicole Francis and Jacques Drouin for helpful comments on the manuscript. We thank the Flow Cytometry, Molecular Biology and Functional Genomics, Disease Modelling and Genome Editing, and Microscopy core facilities at the Montreal Clinical Research Institute (IRCM). We acknowledge the following funding support for this work: M.M. is supported by the Canadian Institutes of Health Research Project Grant 173273; T.M. was supported by the Emmanuel Triassi Foundation and Hommage Jacques Gauthier scholarships; G.D.B. is supported by the Fonds de Recherche Quebec - Santé and SECIHTI scholarships; S.J. and M.U.P. are supported by IRCM foundation scholarships.

## Author contributions

M.M. and T.M. designed the study. T.M. and G.D.B. performed and analyzed the experiments and interpreted the data with help from M.M. E.L. and F.R. assisted with ChIP-seq experiments. J.L. performed mass cytometry data analysis and assisted with parts of RNA-seq analysis. Y.K. performed zygote embryo harvesting and assisted with *ex vivo* embryo culture. S.D., M.U. and S.J. assisted with parts of the experiments. M.M. conceived the project, supervised the study and secured funding. T.M., G.D.B. and M.M. wrote the manuscript with input from all authors. M.M. is the lead contact.

---

**Figure S1.**
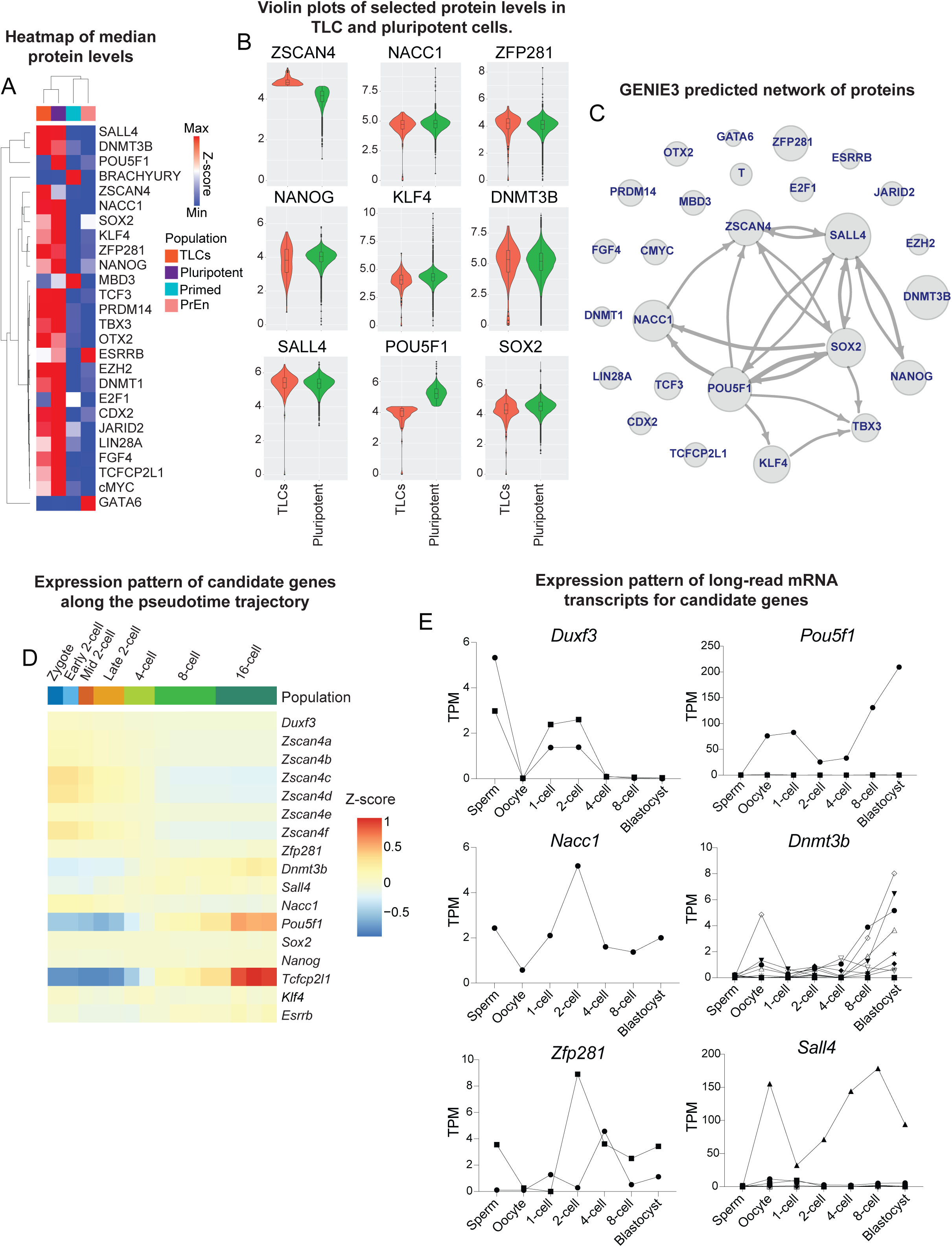
Identification of potential TLC promoting transcription factors. **(A)** Heatmap showing hierarchically clustered median protein levels of transcription factors quantified using mass cytometry in distinct cell states of ESCs: pluripotent, TLCs, primed, and primitive endoderm (PrEn). The protein data is obtained from Meharwade et al. (*62*). **(B)** Violin plots showing the distribution of selected protein levels in TLC and pluripotent state cells. **(C)** GENIE3 predicted network of proteins shown in A. While arrows represent the directionality, their thickness represents the intensity of regulation. Circle sizes denote the levels of the respective protein. **(D)** Expression pattern of candidate transcription factors along the CellRouter predicted pseudotime trajectory from zygote to 16-Cell stage of mouse embryogenesis. **(E)** Quantification of long-read mRNA transcripts for candidate genes (*Duxf3*, *Nacc1*, *Zfp281*, *Pou5f1*, *Dnmt3b and Sall4*) in sperm, oocyte, and preimplantation embryos from zygote to blastocyst stage. Different lines for each gene represent the expression value of its individual transcript isoforms. The long-read transcript data is obtained from Qiao *et al*. (*66*).

**Figure S2.**
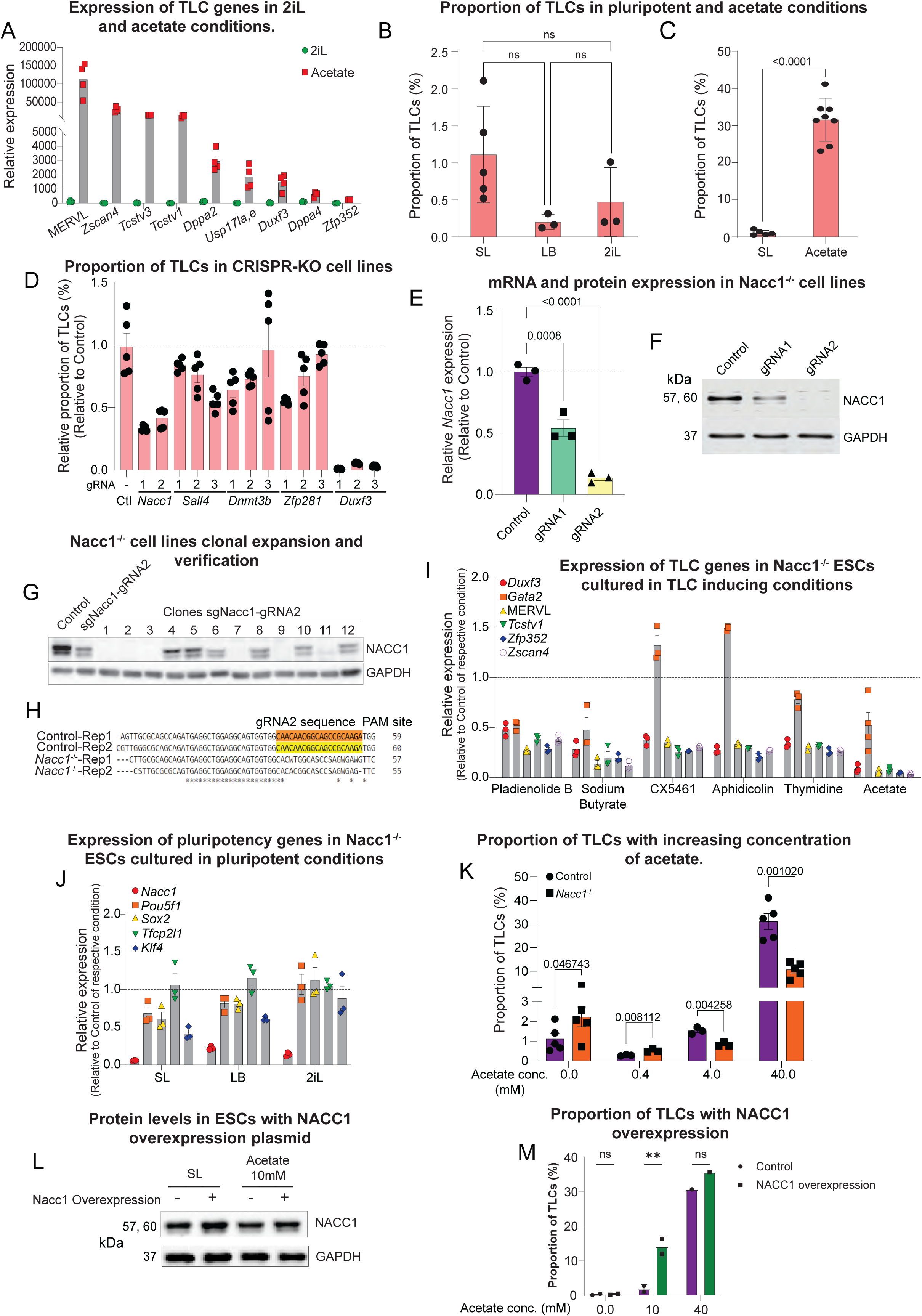
NACC1 induces the TLC state in mouse ESCs. **(A)** RT-qPCR analysis for expression of TLC genes in ESCs cultured in 2iL and acetate conditions. mRNA levels are represented as 2^-ΔΔCt^. **(B)** Proportion of TLCs (MERVL-gag positive cells) in ESCs cultured in SL, LB and 2iL pluripotent conditions. **(C)** Proportion of TLCs in ESCs cultured in SL and acetate conditions. **(D)** Proportion of TLCs in CRISPR-KO cell lines of candidate regulators in the acetate condition. **(E)** Relative expression of *Nacc1* in ESCs with either control, gRNA1 or gRNA2. **(F)** Western blot of NACC1 and GAPDH in ESCs with either control, gRNA1 or gRNA2. **(G)** Western blot for NACC1 and GAPDH in ESCs with control or *Nacc1* gRNA2 (bulk and clones 1-12). **(H)** Sanger sequencing analysis of control and *Nacc1^-/-^* (*Nacc1* gRNA2-Clone2) samples to validate CRISPR-mediated perturbation of *Nacc1* coding sequence near the PAM site. **(I)** RT-qPCR analysis for expression of TLC genes in *Nacc1^-/-^* ESCs cultured in multiple TLC inducing conditions (Methods). **(J)** RT-qPCR analysis for expression of pluripotency genes in *Nacc1^-/-^*ESCs cultured in pluripotency-promoting conditions. **(K)** Proportion of TLCs in control and *Nacc1^-/-^* ESCs cultured with increasing concentration of acetate. **(L)** Western blot for NACC1 and GAPDH in ESCs treated with NACC1 overexpression plasmid in SL and 10mM acetate (Methods). **(M)** Proportion of TLCs in ESCs cultured in SL, 10Mm acetate and 40Mm acetate in control and treated with NACC1 overexpression plasmid. Graphs represent mean ± SEM (in A - E, I-K, M). n=4 (in A), n=3-10 (in B and C), n=5-10 (in D), n=3 (in E), n=3-4 (in I and J), n=3-5 (in K), and n=3 (in M). *P* values were determined by one-way ANOVA in (B and E) and Student’s t-test in (C, K and M).

**Figure S3.**
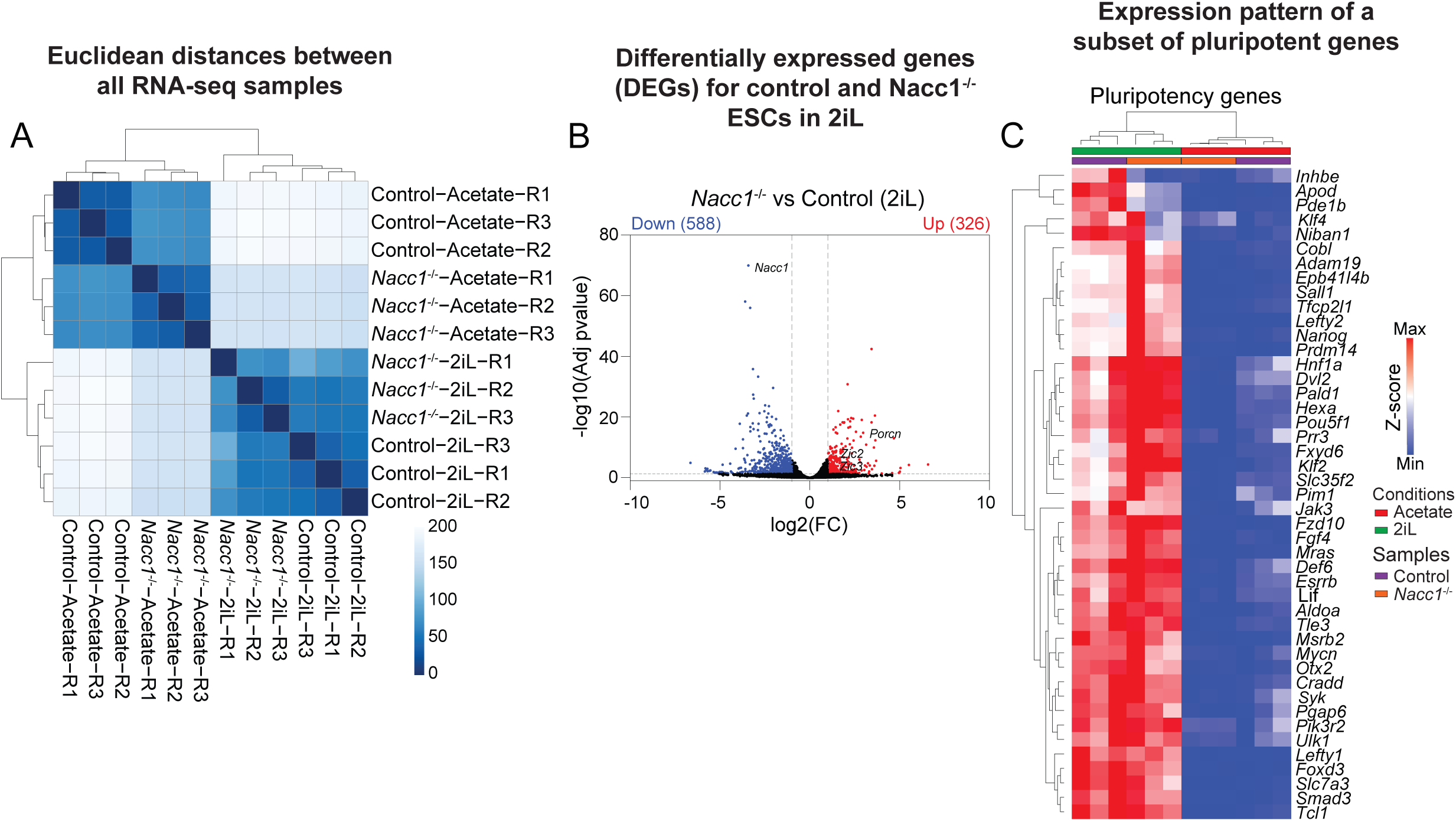
NACC1 knockout has a minor effect on gene expression of ESCs in 2iL culture condition. **(A)** Heatmap of Euclidean distances between all RNA-seq samples based on DESeq2 normalized gene expression. **(B)** Volcano plot showing DEGs for *Nacc1^-/-^* vs control in 2iL condition. Red dots indicate significantly upregulated DEGs, and blue dots indicate significantly downregulated DEGs. Grey line on X-axis indicates log2(FC)±1 and on Y-axis indicates -log10(Adj p-value)=1.301 (lower limit). **(C)** Heatmap showing the differential expression pattern of a subset of pluripotency genes (obtained from Fu *et al.* (*27*)) across control and *Nacc1^-/-^*samples in 2iL or acetate condition comparisons. Columns represent individual samples.

**Figure S4.**
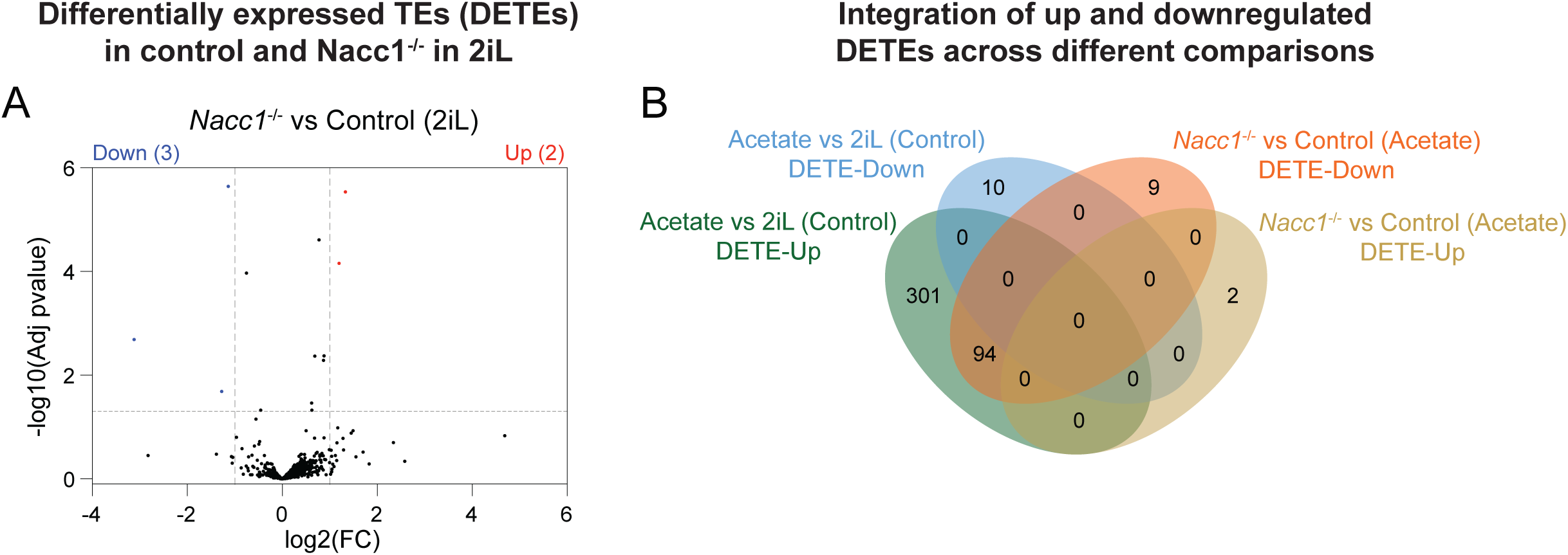
NACC1 knockout has minor effects on the retrotransposons expression in 2iL condition. **(A)** Volcano plot showing DETEs in *Nacc1^-/-^* vs control in 2iL condition. Red dots indicate significantly upregulated TEs and blue dots indicate significant downregulated TEs. Grey line on X-axis indicates log2(FC)±1 and Y-axis indicates -log10(Adj p-value) = 1.301 (lower limit) and 350 (upper limit). **(B)** Venn diagram for the comparison of up and downregulated TEs in acetate vs 2iL (control) and *Nacc1^-/-^* vs control in acetate condition.

**Figure S5.**
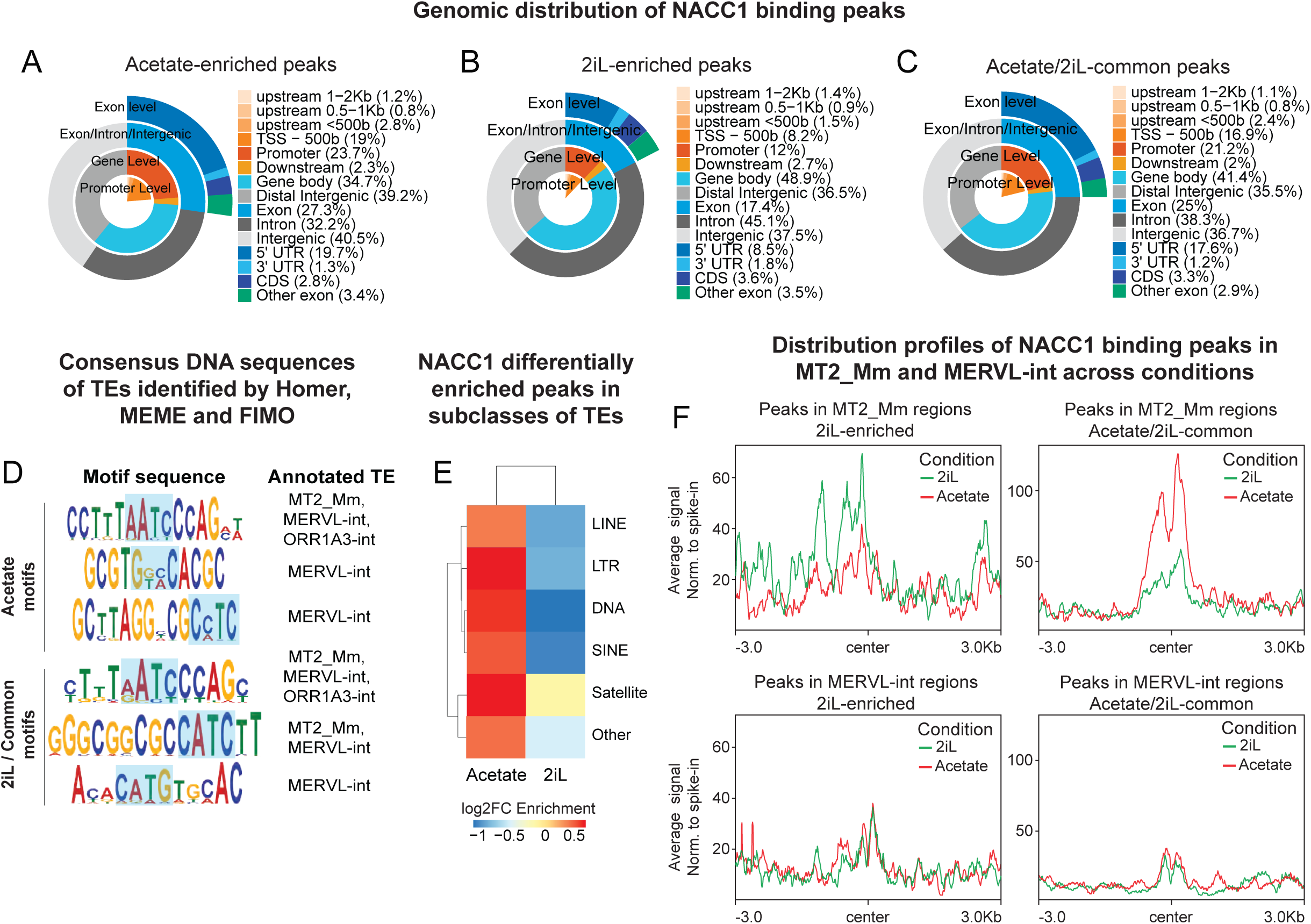
Genomic profile of NACC1 binding in 2iL and acetate conditions. **(A-C)** Genomic distribution of NACC1 bound acetate-enriched (A), 2iL-enriched (B), and acetate/2iL-common (C) peaks. **(D)** Consensus DNA sequences on transposable elements identified by Homer *de novo*, MEME-ChIP, and FIMO motif analysis using NACC1 binding peaks (Methods). The transposable element that is recognized by each sequence is shown. Highlighted region represents the consensus core motif (CATC/G) of recognition for NACC1’s BEN domain. **(E)** Distribution profiles of NACC1 bound peaks at all genomic TEs in 2iL and acetate conditions. **(F)** Heatmap showing NACC1 bound differentially enriched peaks (DEPs) at the indicated subclasses of TEs in 2iL and acetate conditions. **(G)** Distribution profiles of NACC1 2iL enriched and acetate/2iL common peaks at MT2_Mm and MERVL-int retrotransposons in 2iL and acetate conditions.

**Figure S6.**
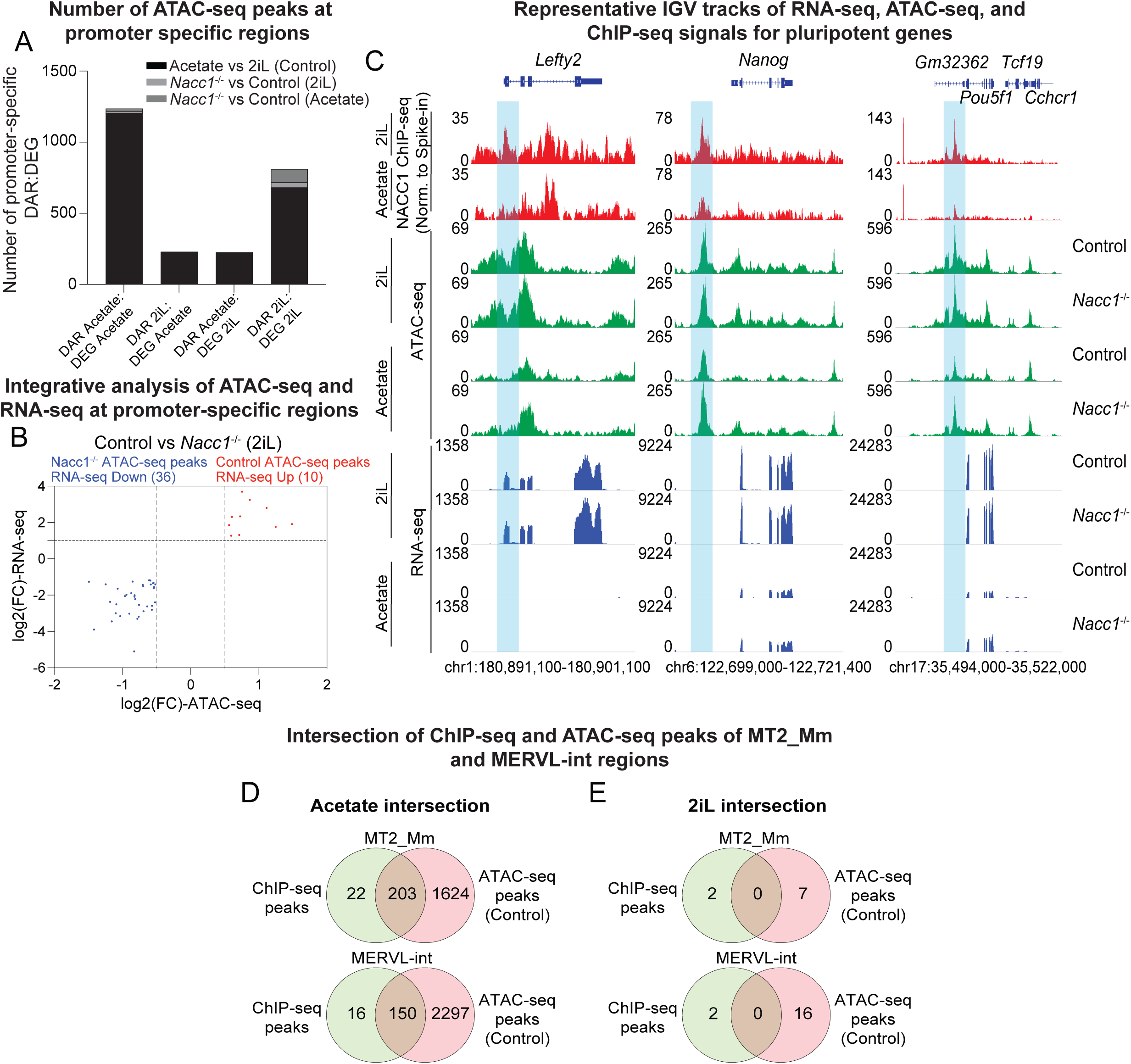
Integration of RNA-seq, ATAC-seq and ChIP-seq datasets. **(A)** Bar plot showing the number of DARs at promoter-specific regions in acetate vs 2iL (control), *Nacc1^-/-^* vs control in 2iL, and *Nacc1^-/-^* vs control in acetate condition. **(B)** Integrative analysis of DARs and DEGs at promoter-specific regions in *Nacc1^-/-^* vs control in 2iL condition. Red dots indicate significantly upregulated genes with open chromatin regions, and blue dots indicate significantly downregulated genes with closed chromatin regions. Grey lines indicate log2(FC)±0.5 on X-axis and log2(FC)±1 on Y-axis. **(C)** Representative IGV snapshots of RNA-seq, ATAC-seq and ChIP-seq signals for pluripotency genes (*Lefty2, Nanog*, and *Pou5f1*). Boxes indicate changes in the peaks in these genes (blue). **(D-E)** Venn diagrams showing the intersection of ChIP-seq and ATAC-seq peaks for MT2_Mm and MERVL-int regions in acetate (D) or 2iL (E).

**Figure S7.**
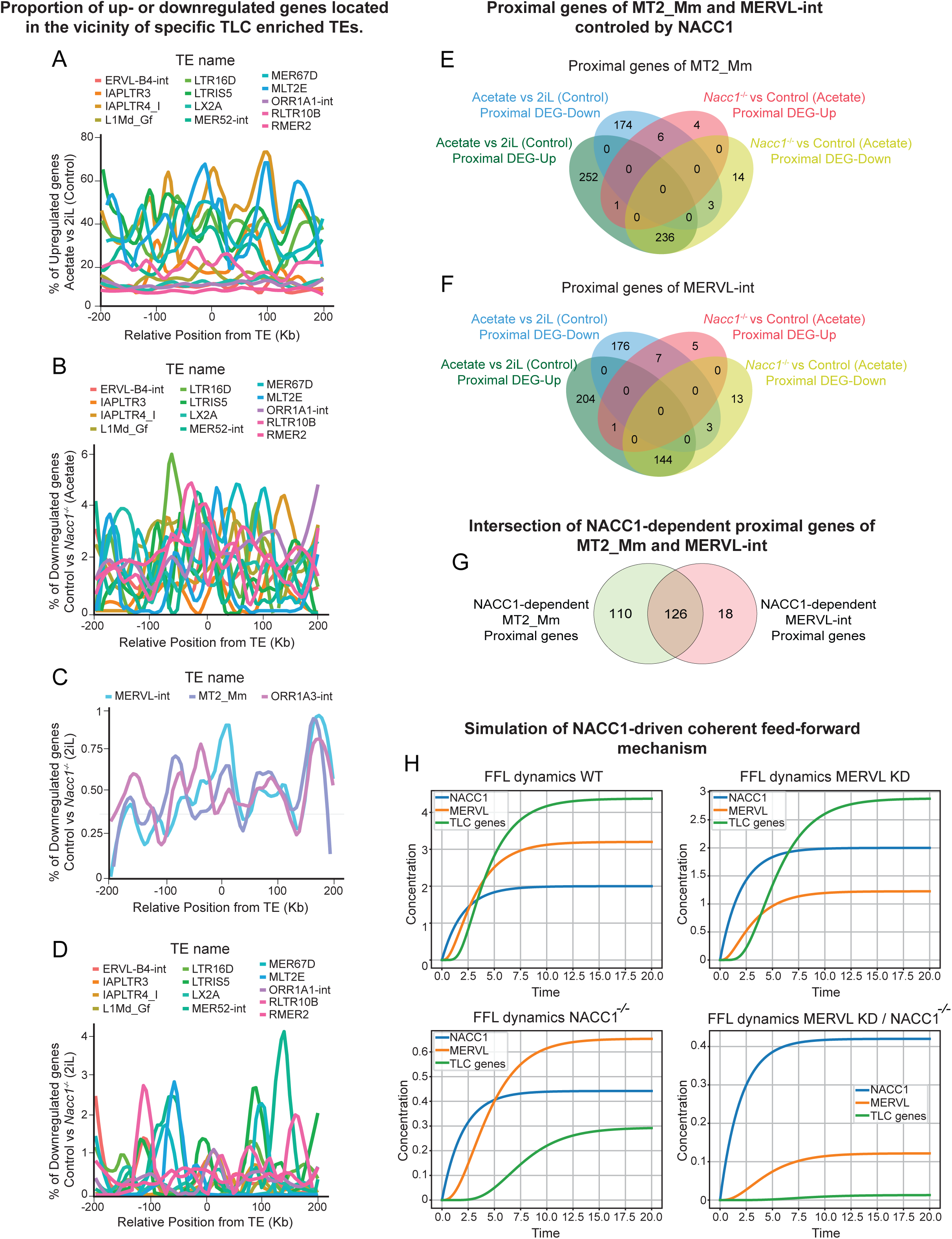
Proximal regulation of genes by TLC associated retrotransposons. **(A-D)** Proportion of up or downregulated genes located in the vicinity of specific TLC enriched TEs. Colored lines represent the percentage trend of upregulated (acetate vs 2iL (control)) (A); downregulated genes (*Nacc1^-/-^*vs control in acetate) (B); and downregulated genes (*Nacc1^-/-^*vs control in 2iL) (C-D). Blue, MERVL-int; purple, MT2_Mm; pink, ORR1A3-int; Red, ERVL-B4-int; orange, IAPLTR3; Goldenrod, IAPLTR4_I; olive green, L1Md_Gf; yellow-green, LTR16D; bright green, LTRIS5; jade green, LX2A; teal green, MER52-int; turquoise, MER67D; blue, MLT2E; purple, ORR1A1-int; hot pink, RLTR10B; salmon pink, RMER2. **(E-F)** Venn diagram showing the comparison between up and downregulated proximal DEGs of MT2_Mm (E) and MERVL-int (F) in acetate vs 2iL (control) and *Nacc1^-/-^* vs control in acetate. **(G)** Venn diagram showing the intersection of NACC1-dependent proximal genes of MT2_Mm and MERVL-int from comparisons in E and F. **(H)** Predicted steady-state dynamics of NACC1 (blue), MERVL (orange), and TLC genes (green) in wild type or knockout/knockdown conditions using an ODE simulation.

## RESOURCE AVAILABILITY

### Lead Contact

Further information and requests for reagents may be directed to and will be fulfilled by the Lead Contact, Mohan Malleshaiah.

### Materials Availability

This study did not generate new unique reagents.

### Data and Code Availability

The high-throughput data reported in this study are available on Gene Expression Omnibus.

## MATERIALS AND METHODS

### Experimental model and subject details Cell culture

Mouse ESCs R1 (E14-Tg2A) and ESCs constitutively and stably expressing EGFP under the EF1α promoter (ref) were routinely cultured by standard methods (*12*). Cells were cultured in knockout DMEM supplemented with non-essential amino acids, sodium pyruvate, L-Glutamine, β- mercaptoethanol, 15% fetal calf serum (Hyclone) and Leukemia inhibitory factor (LIF) (1,000 units/ml, Millipore ESG1106). Cell medium was replaced daily, and cells were passaged every two to three days.

### Conditions to manipulate stem cell fates

ES cells cultured in serum with LIF were dissociated, seeded in the indicated conditions, cultured for 72 h with a daily change of medium, then collected for analysis (gene expression, western blot, RNA-seq, ATAC-seq, ChIP-seq or flow cytometry). For totipotent related conditions, cells were treated with either of the following compounds for the respective durations: Potassium acetate (0.4mM-40mM, 72h, BioBasic, Cat. PRB0438); CX5461 (2μM, 18h, Cayman, Cat. 18392); Aphidicolin (6μM, 18h, Cayman, Cat.14007), Thymidine (2.5mM, 12h treatment followed by release in sL for 9h followed by treatment for 14h, Cayman, Cat.20519), Sodium Butyrate (1mM, 72h, Cayman, Cat. 13121); Pladienolide B (5nM, 72h, Cayman, Cat. 16538). For LBPXRS treatments, cells were cultured in N2B27 containing LIF (1,000 units/ml, Millipore ESG1106), BMP4 (10 ng/ml, Miltenyi Biotech 130-111-168), PD0325901 (1 μM, Millipore Sigma 444968), XAV939 (10 μM, Millipore Sigma 575545), TAK1 inhibitor (1 μM, Tocris 3604), SB431542 (2 μM, Millipore Sigma 616464). For pluripotent related conditions: LB and 2iL, cells were cultured in N2B27 containing either LIF (1,000 units/ml, Millipore ESG1106) and BMP4 (10 ng/ml, Stemgent 03-0007) or LIF (1,000 units/ml, Millipore ESG1106), CHIR99021 (3 μM, Stemgent 04-0004-02) and PD0325901 (1 μM, Millipore Sigma 444968) respectively. All compounds were reconstituted and used as recommended by the manufacturer.

### Generation of OCT4-eGFP:MERVL-gag:tdTomato reporter ESC cell lines

For generation of OCT4-eGFP:MERVL-gag:tdTomato reporter ESC cell lines, OCT-eGFP cell line was seeded in gelatinized dishes and were transfected with MERVL-gag:tdTomato reporter vector (Addgene Plasmid: 40281) using Lipofectamine Stem transfection reagent (ThermoFisher Scientific, STEM00003). After 48hours of transfection, cells were selected with hygromycin for 7 to 10 days. The OCT4-eGFP:MERVL-gag:tdTomato reporter cells were validated by flow cytometry and Leica SP8 laser scanning confocal microscope.

### CRISPR knockout in ESCs

Candidate specific-KO ESC lines were generated by CRISPR knockout technique. Briefly, ESCs were transfected with a blasticidin-resistant lentiCRISPRv2 vector (Addgene Plasmid: 98293) with a guide RNA targeting the exon of candidate genes. After transfection (ThermoFisher Scientific, STEM00003) the cells were blasticidin selected for 7 days and analyzed as bulk-population using flow cytometry. *Nacc1*-KO gRNA2 cells were seeded as single clones (in 0.2μM filtered media containing 1:1 ratio of sL fresh and conditioned media (from wildtype cells)), which were then picked up and expanded. *Nacc1*- KO clones were examined for protein expression by western blotting analysis, and the KO clones were further validated by Sanger sequencing.

### Plasmid DNA Constructs and Transfection

To generate the NACC1 overexpression vector, we first amplified the protein-coding sequence of NACC1 by PCR (primers: forward, 5′- CTCGAGATGGCGCAGACCTTACAGATGGAGATTCCAA −3′; reverse, 5′- GGATCCCTGGAGAACCTCAGCTGAAGGGCCAGCCTC −3′) adding NheI and BamhI sites, and cloned the PCR product into the pmCherry-N1 expression vector (632523, Clontech Takara Bio). Plasmids where verified by nanopore sequencing using Plasmidsaurus. ESCs constitutively and stably expressing EGFP under the EF1α promoter were seeded in gelatinized dishes in SL for 24 hrs. The day after, cells were transfected with PMC-NACC1 plasmid using Lipofectamine Stem transfection reagent (ThermoFisher Scientific, STEM00003) and the media was changed to pluripotent or acetate conditions. After 48hrs of transfection, cells were maintained for 24hrs more in their culture condition and validated by western blot flow cytometry (below).

### Flow cytometry

Single cell suspensions of mESC were evaluated for fluorescence on the BD LSR II, BD LSR Fortessa flow cytometers or were sorted by BDAria (SORP) flow cytometry system at the Montreal Clinical Research Institute platform. Flow cytometry data were analyzed using FlowJo (v10.10.0) (BD Life Sciences).

### Western blotting

Cells were rinsed with PBS and lysed in Laemmli 2X buffer (4% SDS, 20% glycerol, 120 mM Tris-HCl pH 6.8). Extracts were boiled for 10 min, and the protein concentration was assessed with NanoDrop (ThermoFisher). Samples were supplemented with β-mercaptoethanol and bromophenol blue. Extracts were loaded onto SDS-PAGE gels and then transferred to polyvinylidene difluoride (PVDF) membranes. Membranes were incubated with the following primary antibodies: anti-GAPDH (6C5) (Invitrogen AM4300) lot 00755156, and anti-NACC1 (Abcam 29047). Membranes were imaged using ImageQuant LAS 4000 (GE Healthcare). Bands were quantified from immunoblot pictures using Fiji (ImageJ^(ref)^ v1.54g). Signal intensities were normalized over GAPDH intensity.

### Quantitative real-time PCR

Total RNA was purified from samples using the standard TRIzol method. Complementary DNA (cDNA) was prepared using the 5X Prime Script RT Master Mix (Takara RR-036A-1) with 1 μg of total RNA. Reverse transcription was carried out with the standard cycling condition as per the manufacturer’s instructions. cDNA was diluted 30-fold, and quantitative PCR (qPCR) was performed using the PowerUp SYBR Green Master Mix (Thermo Fisher Scientific 100029283). The relative gene expression fold change was calculated using the ΔΔCt method and plotted based on the log-fold expression.

### Generation of blastoids

Cells constitutively and stably expressing EGFP under the EF1α promoter together with MERVL-gag:tdTomato reporter and *Nacc1*-KO gRNA2 cells were cultured in LBPXRS, cell suspension was filtered through a 40 μm cell strainer and then sorted by flow cytometry to obtain totipotent cells (tdTomato+ EGFP+ or tdTomato+ respectively) (*12*). The previously described protocol was used for blastoid generation (*12*). AggreWell 400 (STEMCELL Technologies, 34415) was prepared following the manufacturer’s instructions. ETS-embryo media is composed of 43.8% RPMI 1640 (11875–093), 21.9% DMEM/F-12 (11330–032), and 21.9% Neurobasal (21103–049) media supplemented with 10% FBS (10439–024), 2 mM L-Glutamine (25030–081), 0.5 mM Sodium pyruvate (11360–070), 0.5x of NEAA (11140–050), 0.25x N2 supplement (A13707-01), 0.25x B27 supplement (17504–044), and 0.1 mM 2-mercaptoethanol (21985–023), all from Thermo Fisher Scientific. Blastoid media is further composed of 50% ETS-embryo media and 50% EmbryoMax Advanced KSOM (Sigma, MR-101-D), supplemented with 0.15% BSA (VWR Life Science, 0332), 12.5 ng/mL rhFGF4 (Miltenyi Biotec, 130109388), 0.5 μg/mL Heparin (Sigma-Aldrich, H3149), 0.5 μM A83-01 (Cayman Chemical, 9001799), 5 ng/mL BMP4 (Miltenyi Biotec, 130-110-921), 3 μM GSK3 inhibitor CHIR99021 (Cayman Chemical, 13122), and 2 μM ROCK inhibitor Y-27632 (Cayman Chemical, 10005583). Approximately 7,000 cells (∼5–10 cells per microwell) were resuspended in Blastoid media treated either with DMSO or 20 μM *Nacc1* inhibitor (NIC3) (MedChemExpress, HY-128577) and seeded into one well of the 24- well AggreWell plate. The plate was centrifuged at 200 g for 5 min and transferred into a hypoxia incubator (5% O2, 5% CO2, 90% N2) for 7 days. The day of cell seeding was counted as day 0 of the process. The medium was removed 24 h later (day 1) and replaced with fresh blastoid media containing either DMSO or or 20 μM *Nacc1* inhibitor (NIC3) without Y-27632. Further blastoid media changes were performed at day 4 and day 6. The blastoids were manually picked on day 7 using a mouth pipette (Sigma-Aldrich, A5177) under a stereomicroscope for analysis or downstream experiments.

### Treatment of embryos with NIC3 inhibitors

3-6 weeks old C57BL/6J female mice were superovulated by intraperitoneal injection of PMSG (5IU) followed by 46 hours later hCG (5IU).. After the hCG injection, females were housed overnight with C57BL/6J male mice. Fertilization was assumed to occur approximately 12 hours after hCG injection. Zygotes were collected 17 hours post-hCG injection (corresponding to 5 hours post-fertilization, hpf), screened for the presence of second polar body and transferred to advanced potassium-supplemented simplex optimized medium (KSOM) containing either DMSO (vehicle control) or 20 µM NIC3 at 6 hpf. Experiments described were approved by the Animal Care Committee of the Institut de Recherches Cliniques de Montréal (protocol 2021-09 YK).

### Embryonic development assay

Embryos were cultured in 500 µl of advanced KSOM using a hypoxia incubator (5% O2, 5% CO2, 90% N2). Embryonic development was monitored once every 24 hours using a stereomicroscope (namely 36 hpf, 60 hpf, 84 hpf and 108 hpf). For inhibitor treatment, zygotes were isolated at 5 hours post-fertilization and cultured in inhibitor or mock control from 6 hours post-fertilization onwards. The data was depicted in bar graphs using GraphPad Prism v10.

### Embryo fixation and staining

Embryos were fixed in freshly prepared 4% PFA in PBS for 15 min at room temperature (RT). Fixed embryos were washed three times in PBS without Mg2+/Ca2+ (PBS) containing 0.1% Tween-20 (PBST). Embryos were permeabilized in PBS containing 0.5% Triton X-100 for 30 min (RT), washed 3 times in PBS and incubated in blocking buffer (PBST containing 1% bovine serum albumin, 10% normal donkey serum, and 0.3% Triton X-100) for 1h (RT). Embryos were incubated overnight (4°C) in antibody dilution buffer (ADB) (PBS containing 1% bovine serum albumin, 1% normal donkey serum, and 0.2% Triton X-100) with anti-NACC1 (1:300, Abcam 29047) and MERVL-gag primary antibody (1:100, Novus Biologicals NBP2-66963). Subsequently, they were washed in PBST (3 x 5 min) and incubated with secondary antibody (donkey anti-rabbit AF488 and AF647, 1:1,000 in ADB) at RT for 2hrs in the dark. Next, embryos were washed in PBST (3 x 5 min) and stained with DAPI solution (PBS containing a final concentration of 14.3mM of DAPI). Lastly, embryos were washed in PBST (3 x 5 min) and were transferred into a glass coverslip with 9mm spacers (70327-8S, Electron Microscopy Sciences) under Vectashield (H-1000, Vector Laboratories) mounting medium for imaging. Images were taken on Zeiss LSM710 confocal laser microscope.

### RNA-seq experimental protocol

Triplicates of control and *Nacc1*^-/-^ cells grown in 2iL or Potassium Acetate for 72 h were collected and processed for bulk RNA sequencing. Total RNA was purified from samples using the standard TRIzol method. RNA libraries were prepared from 1000ng of total RNA. Ribosomal RNAs were depleted using the KAPA RiboErase kit (HMR) (Roche Diagnostics; cat num: 08098131702) and libraries were prepared with KAPA RNA Hyperprep Kit (Roche Diagnostics; cat num: 08098107702). Library size distribution was assessed on a 2100 bioanalyzer (Agilent Technologies) and libraries were quantified by qPCR. Equimolar libraries were sequenced in paired-end reads (PE100), on a Novaseq 6000 system (Illumina), with a S4 flowcell and a average coverage of 80M fragments per library.

### RNA-seq computational analysis

The quality of the raw reads was assessed with FASTQC (v0.11.8) (*112*). After examining the quality of the raw reads, trimming was performed with Cutadapt (v4.9) (*113*). Next, we generated a personalized mouse genome that incorporates gene annotation (mm10) with the repeat elements in the mouse genome (rmsk) obtained from UCSC. Next, reads were mapped to this personalized genome using the STAR aligner (v2.7) (*114*), with the options:—readFilesCommand zcat —outFilterMultimapNmax 100 —winAnchorMultimapNmax 100 —outMultimapperOrder Random —runRNGseed 777 — outSAMmultNmax 1 —outSAMtype BAM —outFilterType BySJout —alignSJoverhangMin 8 — alignSJDBoverhangMin 1 —outFilterMismatchNmax 999 —alignIntronMin 20 —alignIntronMax 1000000 —alignMatesGapMax 1000000. Reads mapping to genes and TEs were counted using TEtranscripts (v2.2.3) (*115*) with the option:—mode multi using GTF files for gene annotation downloaded from GENCODE (M12) and GTF files for TE annotations downloaded from the TEtranscripts website (mm10_rmsk_TE.gtf.gz). Differential expression was called using DESeq2 (v1.30.0) (*116*), with FDR < 0.05 (Benjamini-Hochberg-corrected) and an absolute fold-change > 1. Gene ontology analysis was performed using DAVID.

### TE distance analysis

To calculate the distance between genes and repeat elements, the repeat annotation file from RepeatMasker was used for comparison against gene annotation. Custom-made scripts were used to count the total number of genes found in 500-bp bins located at increasing distances from each repeat occurrence or directly overlapping them as previously described (*29*). Custom-made scripts in R (v.4.3.1) were used to obtain the distance from each gene to the closest repeat occurrence for the analyzed retrotransposons. The distance was calculated as the number of base pairs between the annotated transcription start site of a gene to the closest end of a repeat without considering the orientation of the repeat element.

### ATAC-seq experimental protocols

ATAC-seq was performed as previously described and in duplicate for each condition (*117*). A total of ∼50,000 cells were washed once with 50 μl of cold PBS and resuspended in 50 μl ATAC-resuspension buffer (RSB) (10 mM Tris-HCl pH 7.4, 10 mM NaCl, 3 mM MgCl2, 0.1% (v/v) Tween20) supplemented with 0.1% (v/v) IGEPAL CA-630, 0.01% (v/v) Digitonin. The suspension of nuclei was washed once with RSB buffer. Pellets of 50,000 nuclei were tagmented for 30 minutes at 37°C (TDE1 Transposase, Illumina). Tagmented DNA was purified on column DNA clean-5 (Zymo Research). Illumina UDP Indexes were integrated by PCR-enrichment (12 cycles) and final libraries were purified and size selected (L=1.1; R=0,6) with ampure beads (KAPA, Roche Diagnotics). Library size distribution was assessed on a 2100 bioanalyzer (Agilent Technologies) and libraries were quantified by qPCR. Equimolar libraries were sequenced in paired-end reads (PE100), on a Novaseq 6000 system (Illumina), with a S4 flowcell and a coverage of 50M fragments per library.

### ATAC-seq computational analysis

The quality of the raw reads was assessed with FASTQC (v0.11.8) (*112*). After examining the quality of the raw reads, trimming was performed with TRIMMOMATIC (v0.36) (*118*). The reads were aligned to the mouse reference genome with BOWTIE2 (v2.2.6) with mean of 85 % of reads uniquely mapped (*119*). Alignments were post-processed to remove PCR duplicates (Picard tool v2.4.1) and reads mapping to mitochondrial DNA (samtools v1.8) (*120*). In order to represent the real Tn5 transposase binding sites of 9bp, the coordinates of the reads were shifted by +4bp for the plus strand and by −5bp for the minus strand, using deeptools (v3.0.1) (*121*). The former was also used to remove ENCODE’s blacklisted regions (signal artifact regions) and convert bam files to BEDPE format. MACS2 (v2.2.7.1) was used to identify significant peaks (*122*). Diffbind (v2.10.0) R (v3.6.0) package was used to generate the count matrix of Tn5 insertion site numbers for each consensus peak (peaks that were present in all samples) (*123*). Differentially accessible regions (DARs) were identified between conditions using DESeq2 (v1.30.0) R (v3.6.0) package (*116*). DARs were annotated with their closest feature, and different transcription binding motifs were identified with HOMER (v4.8.0) (*124*). A non-biased motif search was also performed with HOMER (v4.8.0), in order to identify all known motifs and *de novo* motifs in these regions. Deeptools (v3.5.1) was used to generate the heatmap figures (*121*). The TE annotation (rmsk) from UCSC was used with Bedtools (v2.25.0) intersect to check overlapping between peaks and TE regions (*125*). Bioinformatics analysis for ATAC-seq was performed at the Bioinformatics core facility of the Montreal Clinical Research Institute (IRCM).

### ChIP-seq experimental protocols

A ChIP assay was carried out as previously described and in duplicates for each condition (*126*). The treated mouse embryonic stem cells were crosslinked with 1% formaldehyde for 10 min and then stopped by adding 125 mM glycine for 5 min at room temperature. Cells were centrifuged at 700g for 3 min at 4°C and washed twice with cold PBS. Equal ratios of protein A beads (Invitrogen 10001D, Lot: 01137460) and protein G beads (Invitrogen 10003D, Lot: 2749885) (30μl total) were washed twice with cold blocking buffer (1x PBS, 0.5% BSA) and incubated with cold blocking buffer mixed with 5 μg of either NACC1 antibody (Ab29047) or IgG antibody (Sc-3888) and 1 μg of spike-in antibody (Active motif #61686) per IP for 5 hrs at 4°C. Chromatin samples were lysed with lysis buffer (50 mM Tris, pH 8.0, 150 mM NaCl, 1% IGE-PAL CA-630, 0.5% Sodium Deoxycholate, 0.1% SDS,1 mM EDTA, and 1x Roche complete protease inhibitors (Roche, 04693159001)) for 15 min on ice. Exactly 1mL was transferred to a 1mL milliTUBE with AFA fiber (Covaris 520130) and sonicated using sonicated with Covaris E220 using the following settings:

| Condition | Number of cells | Peak incident power (w) | Duty factor (%) | Cycles per burst | Duration (s) |
| --- | --- | --- | --- | --- | --- |
| Acetate | $1 \times 10^7$ | 140 | 10 | 200 | 360 |
| 2iL | $1 \times 10^7$ | 140 | 18 | 200 | 300 |

The sonicated cell lysate was centrifuged at max speed for 20 min at 4°C. The soluble chromatin-containing supernatant was divided in 500µL aliquots (equivalent to 5x10^6^ cells) and completed with lysis buffer up to 600 μl; 1% was saved as input controls (equivalent to 5x10^5^ cells). Antibody-coating beads were washed three times with lysis buffer, 30 μl were added to each chromatin tube and incubated overnight with rotation at 4°C. Sequential washes were performed with ChIP wash buffers (A,B,C,D) (ChIP wash buffer A (0.5% IGE-PAL CA-630, 150 mM KCl, 10 mM Tris, 1 mM EDTA,pH 8.0), ChIP wash buffer B (0.5% Triton X-100, 100 mM NaCl, 10 mM Tris), ChIP wash buffer C (0.5% Triton X- 100, 400 mM NaCl, 10 mM Tris), ChIP wash buffer D (0.5% Triton X-100, 500 mM NaCl, 10 mM Tris)), twice with ChIP wash buffer E (0.5% IGE-PAL CA630, 250 mM LiCl, 10 mM Tris, 1 mM EDTA, pH 8.0) and once with TE buffer (10 mM Tris, 1 mM EDTA,pH 8.0). Decrosslinking was carried out in elution buffer (10 mM Tris, 1 mM EDTA, 1% SDS, pH 8.0) at 65 °C for 16 h. RNase A (10 mg/ml) with glycogen (20 mg/ml) and proteinase K (20 mg/ml) with 10% SDS treatments was performed at 37 °C for 2 hrs each.

DNA samples were purified using phenol/chloroform/isoamyl alcohol (sigma-aldrich P2069-400ml) following manufacturing instructions with the following modifications. For ChIP samples phenol-purifed twice, for input samples, phenol-purified three times. Next, NaCl was added to 200mM final concentration to the final aqueous phase and DNA was precipitated by adding 2.5 volumes of cold 100% EtOH. Samples were centrifuged at max speed for 20 min 4 °C. Supernatant was discarded and a wash was performed with cold 70% EtOH. Samples were centrifuged at max speed for 10 min 4 °C. 50 μl of TE buffer was added and samples were incubated for 30 min at room temperature. DNA samples were analyzed using real-time PCR and prepared for deep sequencing. Size distribution and molarity of immunoprecipitated and input samples were evaluated on a 2100 bioanalyzer (Agilent Technologies).

ChIP-seq libraries were prepared using the KAPA hyperprep library kit (Roche Diagnostics). Normalization of the quantities was done after quantification of the ligation products by qPCR. Libraries were amplified by PCR (10 cycles). Library size distribution was assessed on a 2100 bioanalyzer (Agilent Technologies) and libraries were quantified by qPCR. Equimolar libraries were sequenced in paired-end reads (PE100), on a Novaseq 6000 system (Illumina), with a S4 flowcell and a coverage of 50M fragments per library.

### ChIP-seq computational analysis

The quality of the raw reads was assessed with FASTQC (v0.12.0) (*112*). After examining the quality of the raw reads, trimming was performed with TrimGalore (v0.6.10) (*113*). The reads were aligned to either the mouse reference genome (mm10) or drosophila genome (dm6) with BWA (v0.7.17) (*127*). Alignments were sorted and indexed using Samtools (v1.17) and post-processed to remove PCR duplicates (Picard tool v2.26.3) (*120*). NACC1 peaks were called for each genome using MACS2 (v2.2.7.1) with the following parameters (narrowPeak, genome sizes mm: 1.87e9 or dm: 1.2e8, qvalue 0.05) (*122*). The ENCODE’s blacklisted regions (signal artefact regions) was removed for each genome and spike-in reads were used as a normalization factor that equalizes the signal across samples as previously described (*128*). Replicates were merged, sorted, filtered for non-nuclear chromosomes, and converted to bigwig files using deepTools (v3.5.1) (*121*). The heatmap for NACC1 binding scores associated with regions of interest was generated by computeMatrix and plotHeatmap modules in deepTools (v3.5.1). Differentially occupancy analysis between conditions was performed using DESeq2 (v1.32.0) R (v4.3.1) package (*116*). Peak annotation, feature distribution, and functional enrichment analysis was performed using ChIPseeker (v1.28.3), ChIPpeakAnno (v3.26.4) in R (v4.3.1) packages (*129, 130*). Transcription binding motifs and a non-biased motif search was also performed with HOMER (v4.11) and MEME-ChIP, to identify all known motifs and *de novo* motifs in these regions (*124, 131*).

For transposable elements analysis, the annotation for repeat elements in the mouse genome (rmsk) was obtained from UCSC (https://hgdownload.soe.ucsc.edu/downloads.html). To properly evaluate all the repeat elements across the genome, the ENCODE’s blacklisted regions was not removed for both genomes, and the spike-in reads were used to re-normalize the files. Replicates were merged, sorted, filtered for non-nuclear chromosomes, and converted to bigwig files using deepTools (v3.5.1) (*121*). Bedtools (v2.30.0) intersect was used to check overlapping between peaks and TE regions (*125*). Expression profiles of MT2_Mm and MERVL-int transposable elements were generated by computeMatrix and plotHeatmap modules in deepTools (v3.5.1) (*121*). In order to assess the transcription binding motifs on MERVL-int and MT2_Mm sequences, we used the *de novo* motifs that were not annotated to a known gene and used them as input for FIMO together with the consensus sequence of the repeat element obtained from Dfam Database (*132, 133*).

### Intersection of ChIP-seq, RNA-seq and ATAC-seq

For expression profiles of transposable elements identified by ChIP-seq and have an alteration in ATAC-seq, Bedtools (v2.30.0) intersect was used to check overlapping coordinates between technologies and profiles were generated by computeMatrix and plotHeatmap modules in deepTools (v3.5.1) (*125*). Binding and Expression Target Analysis (BETA) was used to integrate ChIP-seq binding sites with RNA-seq differential gene expression data to infer direct target genes of NACC1 (*134*). For the intersection of the three technologies, gene names from each differential expression analysis were selected using the following criteria: RNA-seq log2FC ±1, FDR < 0.05, ChIP-seq and ATAC-seq log2FC ±0.5, FDR < 0.05 and plotted in R (v4.3.1).

### Computational Model of NACC1 Coherent Feed-Forward Loop

To simulate the coherent feed-forward loop (CFFL), we used the ordinary differential equation (ODE) model implemented in Python 3.11 using *odeint* solver from SciPy. The model consist of three components: NACC1 (a totipotent transcription factor), MERVL (a retrotransposable element) and any totipotent gene which expression is dependent of these two factors. The equations describing the production and degradation of each component were formulated with Hill-type activation, such that NACC1 activates MERVL, and both of them can activate totipotent genes (Fig7H). Model parameters included production rates of each component (*α_NACC1, α_Mervl, and α_TLC_genes*), Hill coefficient (*n*), activation thresholds (*K_{nm}, K_{nt}, K_{mt}*), a shared degradation constant (*d*), and initial and final time until the system reaches a stable steady state. The dynamics are given by the following equations:

1. *dNACC1/dt = α_NACC1– d·NACC1* NACC1 is produced at a constant basal rate *α_NACC1* and decays with rate constant *d*.
2. *dMERVL/dt = α_MERVL·(NACC1^n/(K_{nm}^n + NACC1^n)) – d·MERVL* MERVL production follows a Hill activation by NACC1 (maximal rate *α_MERVL*, activation threshold *K_{nm}*, and cooperativity *n*), minus first-order decay.
3. *dTLC_genes/dt = á_TLC_gened·(NACC1^n/(K_{nt}^n + NACC1^n))·(MERVL^n/(K_{mt}^n + MERVL^n)) – d·TLC_genes* Totipotent gene production requires simultaneous activation by NACC1 and by MERVL (modeled as the product of two Hill functions with thresholds *K_{nt}* and *K_{mt}*, cooperativity *n*) with maximal rate *α_TLC_genes*, minus decay. This equation reflects combinatorial regulation, so that TLC gene expression occurs only when both NACC1 and MERVL are present.

To model the steady state, we set the degradation rate at 0.5 to reflect similar transcript half-lives, the Hill coefficient at 2.0 to represent cooperative transcriptional activation, all thesholds at 1.0, the time modeled wass from 0 to 20, and the default production rates to 1.0, 2.0, and 3.0 for NACC1, MERVL, and TLC genes respectively. The production rates of NACC1 and MERVL were then calibrated to reflect experimental mRNA levels observed in their respective knockout and knockdown conditions as shown in the following table:

| Parameter | Default (D) | WT simulation | Mervl KD simulation | Nacc1 KO simulation | MERVL KD / Nacc1 KO simulation |
| --- | --- | --- | --- | --- | --- |
| $\alpha_{NACC1}$ | 1.0 | 1.0 | 1.0 | 0.221*D | 0.21*D |
| $\alpha_{Mervl}$ | 2.0 | 2.0 | 0.383*D | 2.0 | 0.203*D |
| $\alpha_{TLC\_genes}$ | 3.0 | 3.0 | 3.0 | 3.0 | 3.0 |

### ASO-mediated knockdown of MERVL in ESCs

ASO-mediated knockdown was performed using a previously described protocol with minor modifications (*39*). To validate the efficiency of the ASO-mediated KD, the cells were transfected with 10μM ASO against MERVL or scrambled ASOs using Lipofectamine Stem transfection reagent (ThermoFisher Scientific, STEM00003) and seeded into each well of a 24-well plate containing 500 µl ESC medium (SL) with Acetate. After 48h of transfection, the cells were harvested for quantitative RT-PCR to evaluate the expression of MERVL and proximal totipotent genes.

### Quantification and statistical analysis

Quantitative data are presented as means ± SEM. All experiments were independently performed at least three times. Statistical significance of the experiments was determined using Student’s t-tests or One-way ANOVA or Two-way ANOVA; ∗∗∗∗; p < 0.0001, ∗∗∗; p < 0.001, ∗∗; p < 0.01, ∗; p < 0.05, and ns; not significant. Statistical analysis was performed in GraphPad PRISM 10.

